# Human TRMT1 and TRMT1L paralogs ensure the proper modification state, stability, and function of tRNAs

**DOI:** 10.1101/2024.05.20.594868

**Authors:** Kejia Zhang, Aidan C. Manning, Jenna M. Lentini, Jonathan Howard, Felix Dalwigk, Reza Maroofian, Stephanie Efthymiou, Patricia Chan, Sergei I. Eliseev, Zi Yang, Hayley Chang, Ehsan Ghayoor Karimiani, Behnoosh Bakhshoodeh, Henry Houlden, Stefanie M. Kaiser, Todd M. Lowe, Dragony Fu

**Author notes:** Corresponding author: Dragony Fu. Equal contribution as co-first author (Kejia Zhang, Aidan C. Manning).

## Abstract

The tRNA methyltransferase 1 (TRMT1) enzyme catalyzes m2,2G modification in tRNAs. Intriguingly, vertebrates encode an additional tRNA methyltransferase 1-like (TRMT1L) paralog. Here, we use a comprehensive tRNA sequencing approach to decipher targets of human TRMT1 and TRMT1L. We find that TRMT1 methylates all known tRNAs containing guanosine at position 26 while TRMT1L represents the elusive enzyme catalyzing m2,2G at position 27 in tyrosine tRNAs. Surprisingly, TRMT1L is also necessary for maintaining acp3U modifications in a subset of tRNAs through a process that can be uncoupled from methyltransferase activity. We also demonstrate that tyrosine and serine tRNAs are dependent upon m2,2G modifications for their stability and function in translation. Notably, human patient cells with disease-associated TRMT1 variants exhibit reduced levels of tyrosine and serine tRNAs. These findings uncover unexpected roles for TRMT1 paralogs, decipher functions for m2,2G modifications, and pinpoint tRNAs dysregulated in human disorders caused by tRNA modification deficiency.

## Introduction

Over 100 different types of post-transcriptional modifications have been identified in tRNAs (El Yacoubi *et al*, 2012; Jackman & Alfonzo, 2013; Suzuki, 2021). Several of the modifications have been demonstrated to play critical roles in tRNA folding and biogenesis (Phizicky & Hopper, 2023). Moreover, tRNA modifications have emerged as key modulators of gene expression during organismal development, the cellular stress response, and tumorigenesis (Dedon & Begley, 2022; Delaunay *et al*, 2023; Orellana *et al*, 2022; Zhang *et al*, 2022). In particular, the brain appears to be sensitive to perturbations in tRNA modification, as evidenced by the numerous cognitive disorders linked to tRNA modification enzymes (Burgess & Storkebaum, 2023; Chujo & Tomizawa, 2021; Ramos & Fu, 2019). While the connections between tRNA modification and biological pathways are growing, there are still major gaps in knowledge regarding the enzymes responsible for known and predicted tRNA modifications in humans (de Crecy-Lagard *et al*, 2019)

One of the first tRNA modification enzymes identified is tRNA methyltransferase 1 (Trm1p) from the yeast *Saccharomyces cerevisiae* (Hopper *et al*, 1982). Trm1p catalyzes the dimethylation of guanosine at position 26 in cytosolic and mitochondrial tRNAs to yield the *N*2, *N*2-dimethylguanosine (m2,2G) modification. Crystallographic and simulation studies have shown that the m2,2G modification plays a key role in preventing alternative tRNA conformations through destabilizing specific base-pairing interactions (Bavi *et al*, 2013; Pallan *et al*, 2008; Steinberg & Cedergren, 1995; Urbonavicius *et al*, 2006). Consistent with this function, Trm1-null strains in *Saccharomyces cerevisiae* and *Schizosaccharomyces pombe* yeast exhibit defects in tRNA stability that are exacerbated in combination with deletion of either the Trm4p tRNA methyltransferase or the La RNA binding protein (Copela *et al*, 2006; Dewe *et al*, 2012; Porat *et al*, 2023; Vakiloroayaei *et al*, 2017). These observations suggest that Trm1p-catalyzed m2,2G modifications act in concert with additional tRNA modifications and RNA chaperones to assist in the proper folding and stability of tRNA (Porat *et al*, 2021).

In humans, the *tRNA methyltransferase 1* (*TRMT1*) gene encodes a homolog of yeast Trm1p (Bujnicki *et al*, 2002; Liu & Straby, 2000). Like yeast Trm1p, human TRMT1 is required for the formation of m2,2G or m2G in cytosolic and mitochondrial tRNAs (Dewe *et al*, 2017; Jonkhout *et al*, 2021). TRMT1-deficient cells exhibit decreased proliferation rates, alterations in global protein synthesis, and perturbations in redox homeostasis (Dewe *et al*., 2017). Moreover, the TRMT1-catalyzed modification of mitochondrial pre-tRNA-Ile can be erased by the ALKBH7 demethylase to regulate mitochondrial RNA processing and activity (Zhang *et al*, 2021).

Exome sequencing studies have identified pathogenic variants in *TRMT1* as the cause of certain forms of autosomal-recessive intellectual disability (ID) disorders (Blaesius *et al*, 2018; Davarniya *et al*, 2015; Monies *et al*, 2017; Najmabadi *et al*, 2011). Patient cells with biallelic ID-associated TRMT1 variants exhibit global deficits in m2,2G modification (Jonkhout *et al*., 2021; Zhang *et al*, 2020). Consistent with loss-of-function, the ID-associated TRMT1 variants are defective in tRNA binding and enzymatic activity (Dewe *et al*., 2017; Zhang *et al*., 2020). These studies highlight the critical role of TRMT1-catalyzed tRNA modification in ensuring proper development and cognitive function.

In addition to TRMT1, vertebrate genomes encode a TRMT1 paralog encoded by the *tRNA methyltransferase 1-like* (*TRMT1L*) gene (Dewe *et al*., 2017; Jonkhout *et al*., 2021; Towns & Begley, 2012). Both TRMT1 and TRMT1L contain a class I *S*-adenosyl-methionine (SAM)-binding methyltransferase domain and zinc finger motifs (Figure 1a), but methyltransferase activity has only been shown for human TRMT1 (Liu & Straby, 2000). In addition to the methyltransferase domain, TRMT1 contains a mitochondrial targeting signal at the amino terminus and nuclear localization sequence that are each required for TRMT1 localization to the mitochondria and nucleoplasm of human cells, respectively (Dewe *et al*., 2017; Jonkhout *et al*., 2021). In contrast to the nucleoplasmic and mitochondrial localization of TRMT1, TRMT1L is highly enriched in the nucleolus of human cells (Dewe *et al*., 2017; Jonkhout *et al*., 2021). The partial overlap of TRMT1 and TRMT1L in the nucleus suggests that Trm1 paralogs could have evolved new or shared targets of modification that remain to be discovered.

**Figure 1.**
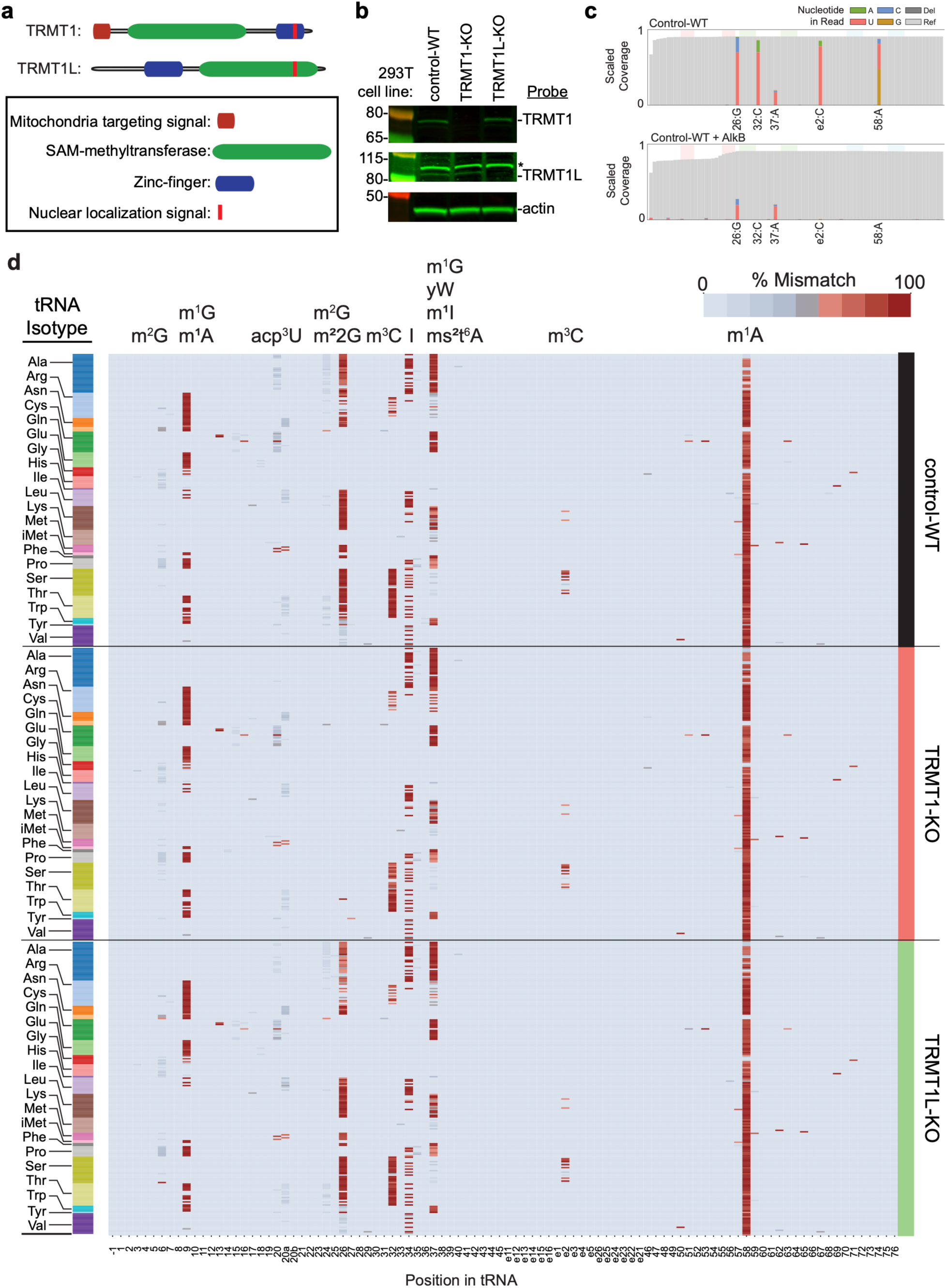
Mapping tRNA modifications dependent upon TRMT1 and TRMT1L. (a) Schematic of TRMT1 and TRMT1L with domains and localization sequences. (b) Immunoblot of TRMT1 and TRMT1L levels in the control-wildtype, TRMT1 and TRMT1L-knockout (KO) cell lines. * denotes a nonspecific band detected by the TRMT1L antibody. (c) Mismatch incorporation frequency across the tRNA transcriptome in control-wildtype cells without (-) or with (+) AlkB treatment. (d) Heatmap of the relative mismatch incorporation frequency across the tRNA transcriptome in control-WT, TRMT1-KO, and TRMT1L-KO cell lines. Predicted modifications at specific positions are noted above the map and nucleotide positions noted below.

Here, we have applied a RNA sequencing approach based upon a retroelement reverse transcriptase to elucidate the human tRNA transcriptome modified by TRMT1 and TRMT1L. This approach has yielded a global overview of tRNA modifications and their dependence on TRMT1 and TRMT1L, including novel roles in tRNA biogenesis and function. Notably, we find that human TRMT1L is an active methyltransferase that catalyzes m2,2G formation in tRNA-Tyr isoacceptors at a position that is unique from TRMT1. These studies reveal molecular roles for m2,2G modifications in tRNA stability and function that are crucial for proper neurodevelopment.

## Results

### Human cytosolic and mitochondrial tRNAs containing G at position 26 are targets of TRMT1

To elucidate the targets of TRMT1 and TRMT1L, we generated human embryonic 293T cell lines deficient in TRMT1 or TRMT1L using CRISPR gene editing. For TRMT1, we made a targeted deletion that removes nearly all of exon 2 containing the start ATG codon in the *TRMT1* gene (Supplemental Figure S1a). For *TRMT1L*, the targeted deletion causes an in-frame deletion in exon 1 containing the start ATG codon (Supplemental Figure S1b). The TRMT1- and TRMT1L-knockout (KO) cell lines were compared to the parental control-wildtype (WT) cell line. We confirmed deletion of the genomic regions in both alleles of the TRMT1- and TRMT1L-KO cell lines using sequencing (Supplemental Data S1). Immunoblotting revealed nearly undetectable TRMT1 or TRMT1L protein in the TRMT1- or TRMT1L-KO cell lines, respectively (Figure 1b).

To monitor changes in the tRNA transcriptome caused by TRMT1 or TRMT1L, we performed Ordered Two-Template Relay (OTTR)-seq, a comprehensive, low-bias sequencing-based methodology based upon a modified retroelement reverse transcriptase (Upton *et al*, 2021). Using the OTTR-seq approach in these cell lines, we generated full-length read coverage for 206 of the 258 total tRNAs in the high-confidence human tRNA set (Chan *et al*, 2021), encompassing all 46 of the canonical tRNA isoacceptor families (Supplemental Figure S2a-c). We also detected all 22 of the human mitochondrial tRNAs.

We next leveraged modification-induced misincorporations in the data to reveal potential changes in base modifications due to the loss of TRMT1 or TRMT1L. Initially, we validated that the misincorporations in our dataset were indicative of known base modifications based upon pre-treatment of samples with the *E. coli* AlkB demethylase that can remove m1A, m3C, and m1G modifications (Clark *et al*, 2016; Cozen *et al*, 2015). We observed a high percentage of misincorporation at position 58 indicative of the m1A modification that was abolished by AlkB treatment (Figure 1c, position 58:A). Moreover, we detected a strong misincorporation signature at position 26 indicative of the m2,2G modification that was reduced but not completely abolished after pre-treatment with AlkB (Figure 1c, position 26:G). These results are consistent with the known demethylase activity of AlkB on these different types of modifications (Dai *et al*, 2017; Zheng *et al*, 2015).

Investigation of modification-induced misincorporations across all cytosolic tRNA transcripts in the control-WT cells revealed additional sites of tRNA modification known to cause RT misincorporations, such as m1G at positions 9 and 37, acp3U at position 20, and m3C at position 32 (Figure 1d, Supplemental Figure S2d). In general, the modification profiles match those expected from previous studies in human cells (Behrens *et al*, 2021; Clark *et al*., 2016; Cui *et al*, 2021; Hernandez-Alias *et al*, 2023; Pinkard *et al*, 2020). Focusing on position 26, we identified significant misincorporations indicative of m2,2G or m2G in 132 of 234 cytosolic tRNAs (Figure 1d, Supplemental Figure S2d, Supplemental Table 1). The misincorporation signature at position 26 for tRNA-Val and tRNA-iMet was distinct from other tRNAs, consistent with the presence of m2G instead of m2,2G in these tRNAs. Notably, we observed the complete loss of misincorporations at position 26 in cytosolic tRNAs of TRMT1-KO cells, but not TRMT1L-KO cells (Figure 1d, Supplemental Figure S3).

In addition to cytosolic tRNAs, we detected a misincorporation signature for m2G or m2,2G modification at position 26 in 5 of 22 mitochondrial tRNAs from control-WT cells (Supplemental Figure S4a, Table 2). The misincorporation signature at position 26 in the mitochondrial tRNAs was abolished in TRMT1-KO cells but still present in the TRMT1L-KO cell line (Supplemental Figure S4b and 4c). While m2G or m2,2G modification at position 26 has been identified in a subset of human mitochondrial tRNAs (Suzuki *et al*, 2020), the cellular requirement for TRMT1 in the modification of mt-tRNA-Glu or mt-tRNA-Asn has not been tested. To validate the role of TRMT1 in the modification of mt-tRNA-Glu and mt-tRNA-Asn, we used a primer extension assay in which the presence of m2G leads to a reverse transcriptase (RT) pause at position 26 (Motorin *et al*, 2007; Youvan & Hearst, 1979). We found that the m2G modification at position 26 in mt-tRNA-Asn and mt-tRNA-Glu was reduced to background levels in the TRMT1-KO cell lines but retained in the TRMT1L-KO cell line (Supplemental Figure 5). Altogether, these studies demonstrate that TRMT1 is the primary enzyme responsible for all detectable m2G or m2,2G modifications at position 26 in nuclear- and mitochondria-encoded tRNAs.

### TRMT1 and TRMT1L are required for distinct m2,2G modifications in tyrosine tRNAs

In contrast to TRMT1, no enzymatic activity has been described for its paralog TRMT1L. As noted above, the misincorporation frequency at position G26 in tRNAs was abolished in TRMT1-KO cells but not significantly impacted in TRMT1L-KO cell lines (Figure 2a, Supplemental Figure S6a). Interestingly though, we observed a reduction in misincorporation frequency at position 27 in tRNA-Tyr isodecoders of the TRMT1L-KO strain, but not TRMT1-KO cells (Figure 2a, Supplemental Figure S6b). Consistent with the reduction in modification at position 27 of tRNA-Tyr, we observed an increase in read coverage for full-length mature tRNA-Tyr-GUA in the TRMT1L-KO cell line caused by increased read-through by the RT (Supplemental Figure S6c). Unlike all other human tRNAs, the tRNA-Tyr isoacceptor family exhibits the unique property of containing two consecutive m2,2G modifications at positions 26 and 27 (Figure 2b) (Johnson *et al*, 1985; Pan, 2018; van Tol *et al*, 1987). These data suggest that TRMT1 modifies position 26 in tRNA-Tyr while TRMT1L represents the enzyme that modifies position 27 in tRNA-Tyr.

**Figure 2.**
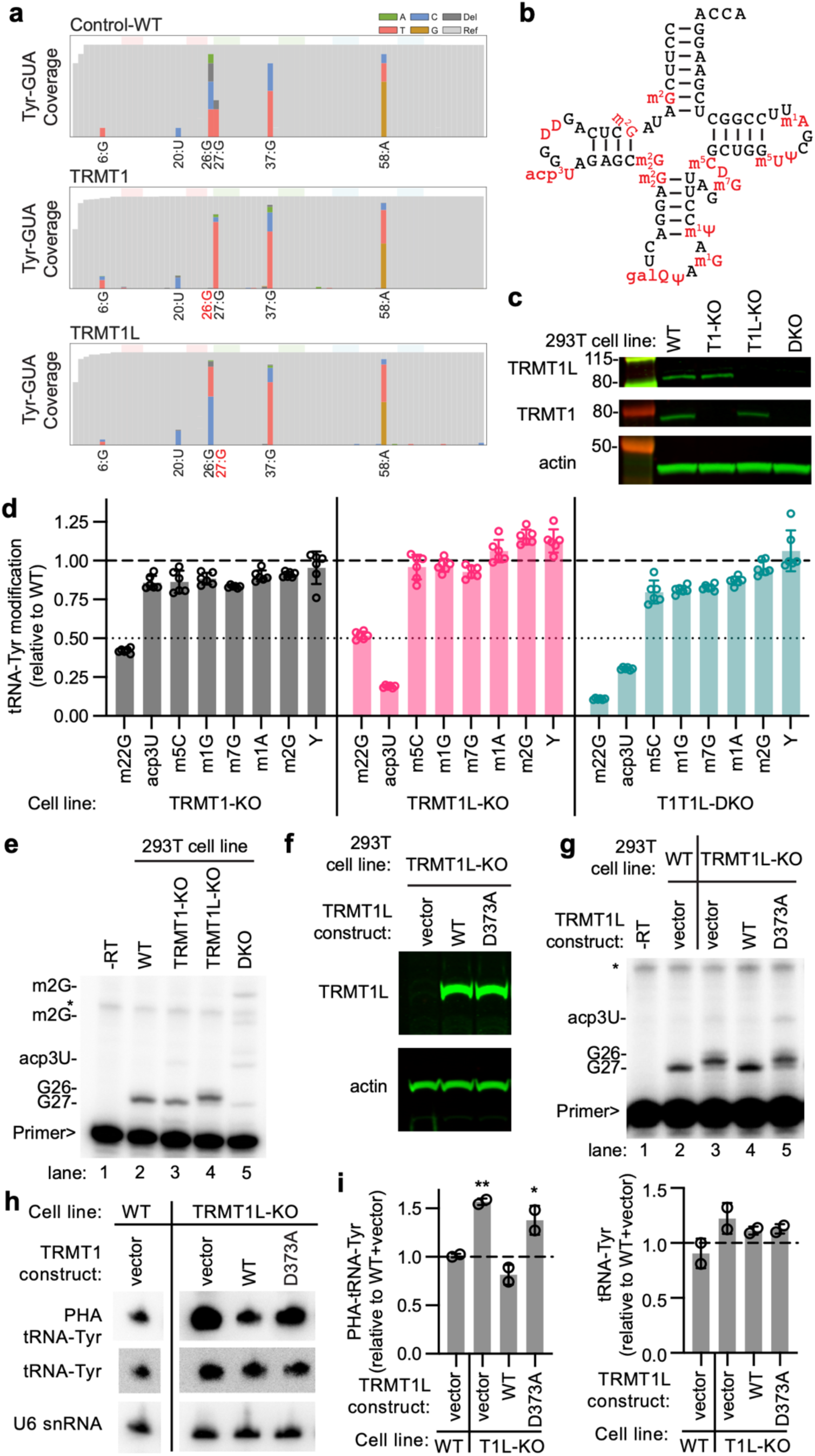
TRMT1 and TRMT1L catalyze m2,2G modification at positions 26 and 27 of tRNA-Tyr, respectively. (a) Misincorporation frequency for tRNA-Tyr isoacceptors from the indicated cell lines. (b) Secondary structure of tRNA-Tyr with modifications noted. (c) Immunoblot of TRMT1 and TRMT1L levels in the control wildtype (WT), TRMT1 (T1) and TRMT1L (T1L)-knockout (KO) cell lines. (d) Relative peak intensity values of the indicated modifications detected by LC-MS in purified tRNA-Tyr from the TRMT1-KO, TRMT1L-KO, and TRMT1-TRMT1L-double knockout (DKO) cell lines versus the control-WT cell line. (e) Primer extension analysis of tRNA-Tyr from the indicated cell lines. * represents a non-specific band present in all samples. (f) Immunoblot analysis confirming expression of TRMT1L variants in the TRMT1L-KO cell line. (g) Primer extension analysis of tRNA-Tyr from control-WT or TRMT1L-KO cell lines with stable integration of the indicated TRMT1L constructs. (h) Northern blot analysis using a PHA probe designed to detect m2,2G at position 26 and a control probe that hybridizes to a different area of the same tRNA for normalization. (i) Quantification of Northern blot signal in (h). PHA quantification represents the ratio of PHA versus control probe signal expressed relative to the control-WT cell line. tRNA levels were normalized using U6 snRNA as a loading control. Bars represent the standard deviation from the mean. Statistical analysis was performed using one-way ANOVA and significance calculated using Dunnett’s multiple comparison test. **P ≤ 0.01. *P ≤ 0.05.

To investigate the role of TRMT1 and TRMT1L in the modification of tRNA-Tyr, we generated a TRMT1-TRMT1L double knockout (DKO) cell line to ascertain whether there was redundancy in either of the modification enzymes. The DKO cell line was generated in a similar manner as above by removing exon 1 of the TRMT1L gene in the TRMT1-KO cell line described above (Supplemental Figure S1). Immunoblotting revealed the absence of detectable TRMT1L protein in both the TRMT1L-KO and DKO cell lines compared to the control-WT cell line (Figure 2c).

To directly test the presence of m2,2G modifications, we isolated tyrosine tRNA isoacceptors using biotin-oligonucleotides followed by nucleoside analysis of the nuclease-digested samples using liquid chromatography-mass spectrometry (LC-MS). The LC-MS approach also allowed us to monitor other modifications that were not detected using the OTTR-Seq approach. We find that tyrosine tRNAs from TRMT1-KO cells contain ∼50% of the m2,2G modification compared to control-WT cell lines (Figure 2d, TRMT1-KO). Tyrosine tRNAs from TRMT1L-KO cells also contain ∼50% of the m2,2G modification compared to control-WT cell lines (Figure 2d, TRMT1L-KO). The reduction in m2,2G modification by approximately half in the TRMT1-KO or TRMT1L-KO cell lines is consistent with each enzyme being responsible for one m2,2G modification in tRNA-Tyr. Furthermore, we find that m2,2G modification in tyrosine tRNAs is reduced to near background levels in the TRMT1-TRMT1L DKO cell line (Figure 2d, T1T1L-DKO). In addition to m2,2G modifications, we found that the 3-(3-amino-3- carboxypropyl)uridine (acp3U) modification in tRNA-Tyr was decreased in the TRMT1L and DKO cell lines (Figure 3d, acp3U).

**Figure 3.**
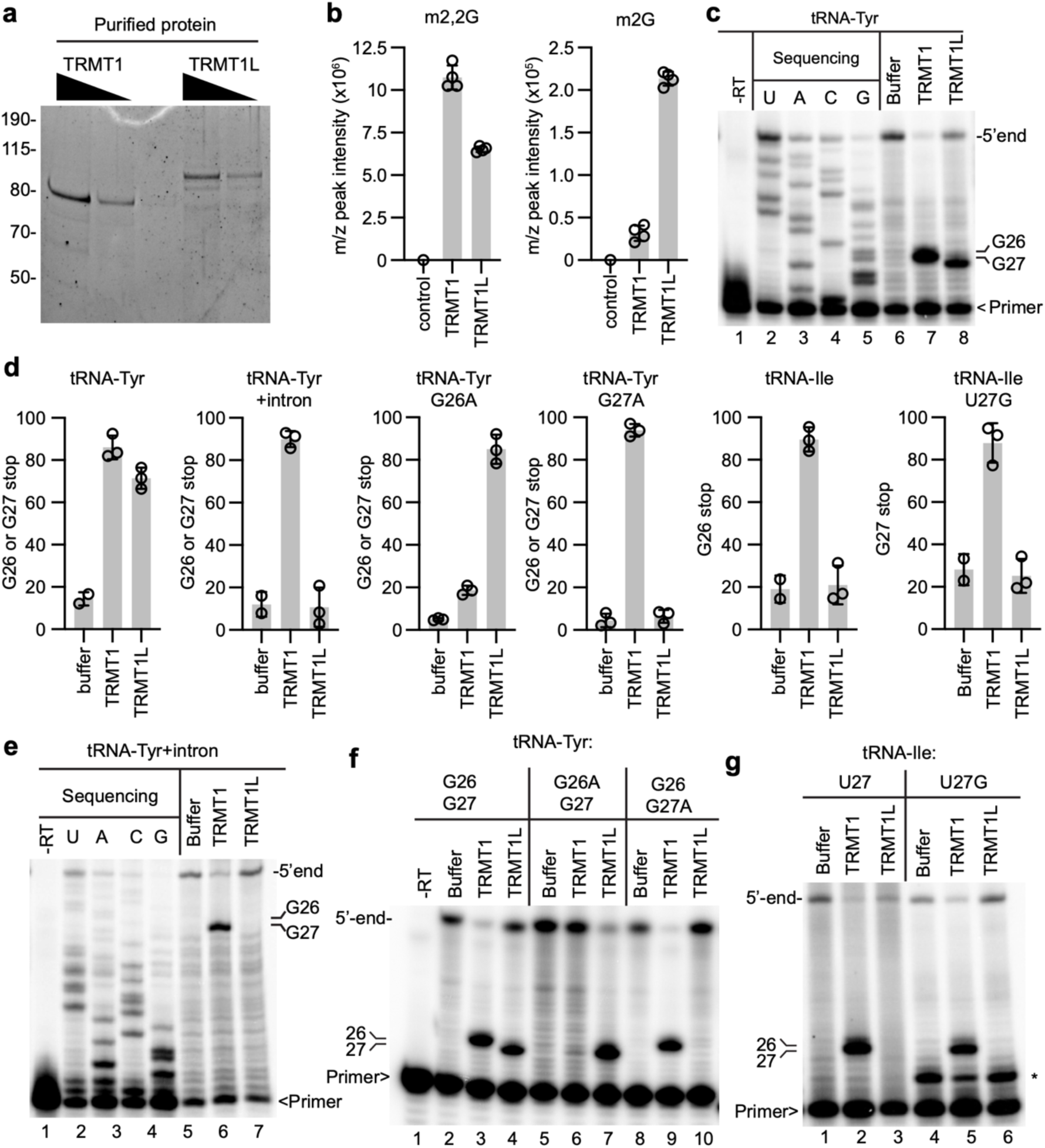
In vitro reconstitution of TRMT1 and TRMT1L methyltransferase activity on tRNA substrates. (a) Sypro Ruby stained gel of purified Strep-TRMT1 and TRMT1L from human 293T cells. (b, c) Peak intensity areas of m2,2G or m2G measured by liquid chromatography-mass spectrometry (LC–MS) in *in vitro* transcribed tRNA-Tyr after pre-incubation with buffer, TRMT1, or TRMT1L. Peak intensity is normalized to the canonical nucleosides A, U, G, and C. (c) Representative gel of primer extension assay to monitor m2,2G in tRNA-Tyr after incubation with buffer, TRMT, or TRMT1L. (d) Quantification of G26 and/or G27 stop in (c), (e), (f), and (g). (e) Primer extension analysis of intron-containing pre-tRNA-Tyr after pre-incubation with the indicated buffer or proteins. (f) Primer extension analysis of the indicated tRNA-Tyr variants after incubation with buffer or the indicated proteins. (g) Primer extension analysis of the tRNA-Ile and tRNA-Ile-U27G after incubation with buffer or the indicated proteins.

We also independently confirmed the changes in tRNA-Tyr modification state in TRMT1-KO, TRMT1L-KO, and DKO cell lines using absolute quantification via mass spectrometry coupled to stable isotope-labeled internal standards (Heiss *et al*, 2021; Kellner *et al*, 2014). Using this approach, we detected two m2,2G modifications per tRNA-Tyr molecule from control-WT human cells (Supplemental Figure S7). Consistent with results above, the amount of m2,2G in tRNA-Tyr decreased to approximately one m2,2G per tRNA-Tyr from the TRMT1-KO or TRMT1L-KO cell lines and decreased to near background levels in the DKO cell line (Supplemental Figure S7).

To validate that TRMT1 and TRMT1L are required for methylating positions 26 and 27, respectively, we developed a primer extension assay to resolve the consecutive m2,2G modifications in tRNA-Tyr isoacceptors at single nucleotide resolution. Using this assay, we could detect a stop at position 27 with minimal readthrough products in control-WT cells while a control reaction without RT yielded no detectable stop at position 27 (Figure 2e, lanes 1 and 2, see Supplemental Figure S8 for sequencing ladder). In the TRMT1-KO cell line, the stop at position 27 was still detectable, consistent with TRMT1L modification (Figure 2e, lane 3). We note that a slight, but reproducible readthrough product could be detected in the TRMT1-KO cell line indicating that position 26 has been affected. In the TRMT1L-KO cell line, the RT block at position 27 was greatly diminished with the appearance of a RT block one nucleotide upstream at position 26 (Figure 2e, lane 4). Notably, the block at position 26 was abolished in the DKO cell line with readthrough products up to the acp3U modification at position 20 and m2G modifications at positions 9 and 6 (Figure 2e, lane 5, Supplemental Figure S8). These results are consistent with TRMT1 and TRMT1L methylating positions 26 and 27, respectively, in tRNA-Tyr isoacceptors.

To confirm the role for TRMT1L in formation of m2,2G at position 27 in tRNA-Tyr, we performed rescue experiments via re-expression of wildtype or mutant versions of TRMT1L in the TRMT1L-KO cell line. We created a construct that expresses a TRMT1L mutant in which the predicted aspartic acid residue that binds *S*-adenosyl-methionine in the methyltransferase catalytic site was changed to alanine (TRMT1L-D373A). We generated TRMT1L-KO cell lines with empty vector or constructs expressing wildtype (WT) TRMT1L or TRMT1L-D373A mutant (Figure 2f). As shown above, tRNA-Tyr from control-WT cells exhibited a primer extension stop at position 27 that was not detected without RT (Figure 2g, lanes 1 and 2). Confirming our results above, the TRMT1L-KO cell lines with vector alone exhibited background levels of m2,2G stop at position 27 in tRNA-Tyr compared to wildtype control cells (Figure 2g, lane 3). Re-expression of wildtype TRMT1L in TRMT1L-KO cells was sufficient to restore m2,2G formation at position 27 in tRNA-Tyr (Figure 2g, lane 4). Only background levels of m2,2G was detected at position 27 upon re-expression of the TRMT1L variant with catalytic site mutations in the TRMT1L-KO cell lines (Figure 2g, lane 5).

As further validation for a role of TRMT1L in tyrosine tRNA modification, we employed the Positive Hybridization in the Absence of modification (PHA) assay (Arimbasseri *et al*, 2015; Khalique *et al*, 2022). This Northern blot-based assay relies on differential probe hybridization to tRNA caused by the presence or absence of m2,2G, which impairs base-pairing. Thus, a decrease in m2,2G modification leads to an increase in PHA probe signal that can be normalized against the probe signal from a different region of the same tRNA as an internal control. Using this approach, we detected an increase in PHA probe signal for tRNA-Tyr-GUA in the TRMT1L-KO cell line compared to control-WT cells (Figure 2h, i). The increased PHA probe hybridization for tRNA-Tyr in TRMT1L-KO cells is consistent with loss of m2,2G at position 27 in tyrosine tRNAs. Re-expression of WT-TRMT1L in the TRMT1L-KO cell lines reduced the PHA signal back to wildtype levels compared to TRMT1L-KO cells with vector (Figure 2h, i, WT). In contrast, expression of the TRMT1L-D373A mutant had no major change in the PHA signal of tRNA-Tyr-GUA in the TRMT1L-KO cell line compared to vector alone (Figure 2h, i, D373A). Altogether, these results indicate that TRMT1 and TRMT1L are required for the m2,2G modification at positions 26 and 27, respectively, in tyrosine tRNAs.

### TRMT1 and TRMT1L catalyze m2,2G formation at specific positions in tRNA-Tyr

While tRNA methyltransferase activity has been demonstrated for human TRMT1 (Liu & Straby, 2000), the enzymatic activity of TRMT1L remains to be shown. Thus, we tested the methyltransferase activity of purified TRMT1 or TRMT1L on *in vitro* transcribed tRNA-Tyr substrates. TRMT1 or TRMT1L fused to the tandem Strep tag were expressed and purified from human cells using the Strep-tag purification system (Figure 3a). We incubated the purified TRMT1 or TRMT1L with tRNA-Tyr in the presence of *S*-adenosyl-methionine and monitored m2,2G or m2G formation using LC-MS as described above. No m2,2G or m2G modification was detected in tRNA-Tyr after pre-incubation with buffer from a mock control purification (Fig. 3b). In contrast, pre-incubation of tRNA-Tyr with purified TRMT1 or TRMT1L revealed the formation of m2,2G as well as m2G (Figure 3b). Under these conditions, TRMT1 catalyzes the methylation of guanosine from m2G to m2,2G in a more efficient manner than TRMT1L since more m2G is detected in the tRNA-Tyr substrate with TRMT1L compared to TRMT1 (Figure 3b).

We next verified that the *in vitro* activity of TRMT1 and TRMT1L was specific to positions 26 and 27 of tRNA-Tyr, respectively, using the primer extension assay noted above. No extension product was detected with tRNA-Tyr in the absence of RT (Figure 3c, lane 1). Primer extension with tRNA-Tyr pre-incubated with buffer led to the generation of full-length product with only background signal at positions 26 and 27 (Figure 3c, lane 6). Pre-incubation of tRNA-Tyr with purified TRMT1, followed by primer extension, revealed the appearance of an RT block indicative of m2,2G formation at position 26, but not position 27 (Figure 3c, lane 7, quantified in 4d). The tRNA-Tyr incubated with TRMT1L exhibited a predominant RT stop at position 27 (Figure 3c, lane 8, quantified in 3d). Thus, purified TRMT1 and TRMT1L can catalyze m2,2G formation in tRNA-Tyr at positions 26 and 27, respectively.

Since all human tRNA-Tyr genes contain an intron, we also tested the activity of TRMT1 and TRMT1L on *in vitro* transcribed pre-tRNA-Tyr substrates with an intron. No extension product was detected in the absence of RT (Figure 3e, lane 1). Primer extension with pre-tRNA-Tyr incubated with buffer led to the generation of full-length product with only background signal at positions 26 and 27 (Figure 3e, lane 5). An RT block was detected at position 26 in pre-tRNA-Tyr after incubation with TRMT1 (Figure 3e, lane 6, quantified in 4d). In contrast, no RT block was detected at either position 26 or 27 in pre-tRNA-Tyr after incubation with TRMT1L (Figure 3e, lane 7, quantified in 3d). These results show that TRMT1 can catalyze m2,2G formation in mature as well as pre-tRNA-Tyr substrates containing an intron while TRMT1L exhibits detectable activity only on mature tRNA-Tyr.

In addition to wildtype tRNA-Tyr substrates, we tested the activity of TRMT1 and TRMT1L on tRNA-Tyr substrates in which G26 or G27 was mutated to A (G26A or G27A). Replicating our results above, pre-incubation of wildtype tRNA-Tyr-GAU containing G26 and G27 with purified TRMT1 or TRMT1L, followed by primer extension, revealed the appearance of an RT block indicative of m2,2G formation at positions 26 or 27, respectively (Figure 3f, lanes 3 and 4). Intriguingly, we can detect a minor stop signal at position 27 instead of 26 with the tRNA-Tyr G26A variant after incubation with TRMT1 (Figure 3f, lane 6, quantified in 4d). We could still detect the formation of m2,2G at position 27 for the tRNA-Tyr G26A variant incubated with TRMT1L (Figure 3f, lane 7, quantified in 3d). This result suggests that TRMT1L can modify G27 independently of whether G26 is modified while TRMT1 can modify G at position 27 when the preceding nucleotide position is no longer G.

For the tRNA-Tyr G27A variant, we could detect an RT block at position 26 after incubation with TRMT1 (Figure 3g, lane 9, quantified in 3d). In contrast, no RT block above background was detected at position 26 or 27 of tRNA-Tyr-G27A after incubation with TRMT1L (Figure 3g, lane 10, quantified in 3d). This result indicates that TRMT1L has specificity for G at position 27 and does not switch to modifying position 26 if position 27 is no longer a modification target.

As another test for the specificity of the enzymes, we tested the activity of TRMT1 and TRMT1L on an *in vitro* transcribed tRNA-Ile-AAU, which contains m2,2G at position 26 *in vivo* as we have shown above (Figure 1). Incubation of tRNA-Ile-AAU with TRMT1, but not TRMT1L, led to the formation of an RT block at position 26 indicative of m2,2G formation (Figure 3g, lanes 2 and 3, quantified in 3d). In addition, we tested whether mutation of the A at position 27 of tRNA-Ile-UAU to G could convert tRNA-Ile-AAU to a TRMT1L substrate. While an RT block at position 26 was detected in tRNA-Ile A27G after incubation with TRMT1, only background signal was detected at positions 26 or 27 after incubation with TRMT1L (Figure 3g, lanes 5 and 6, quantified in 3d). These results suggest that TRMT1L recognizes elements distinct to tRNA-Tyr in order to methylate position 27.

### TRMT1 or TRMT1L-deficiency impacts acp3U modification status in a distinct subset of tRNAs

In addition to m2,2G modification at position 26 and 27, we compared the misincorporation frequency at all other positions across the tRNA transcriptome between wildtype, TRMT1-KO, and TRMT1L-KO cell lines (Supplemental Figure S3). A change in misincorporation frequency was detected at position 20 of tRNA-Ala isodecoders, which was reduced in the TRMT1-KO cell lines compared to control-WT cells (Figure 4a, Ala). As shown above, tRNA-Ala contains m2,2G at position 26 catalyzed by TRMT1. Surprisingly, we also found that TRMT1L-KO cell lines exhibited a reduction in misincorporation at position 20 in tRNA-Ala as well as tRNA-Cys even though neither of these tRNAs contain m2,2G modification formed by TRMT1L (Figure 4a, Ala and Cys). We note that TRMT1L-KO cells also exhibit a reduction in acp3U modification in tRNA-Tyr as shown above (Figure 2d). Most tRNAs contain dihydrouridine at position 20 in the D-loop. However, dihydrouridine does not lead to modification-induced misincorporations using this methodology based on our observations, as well as previous findings (Clark *et al*., 2016). Instead, the misincorporation pattern is consistent with the acp3U modification that is known to be present in human tRNA-Tyr, -Ala, -Cys, -Asn, and -Ile isoacceptors (Behrens *et al*., 2021; Cappannini *et al*, 2023; Takakura *et al*, 2019). In human cells, the DTWD1 enzyme catalyzes the majority of acp3U modification in tRNA-Cys and tRNA-Tyr while the DTWD2 enzyme catalyzes the formation of acp3U in tRNA-Asn and tRNA-Ile isoacceptors (Takakura *et al*., 2019). The D-loop of tRNA-Ala isoacceptors matches the D-loop in tRNA-Cys and tRNA-Tyr that is thought to be the recognition sequence of DTWD1.

**Figure 4.**
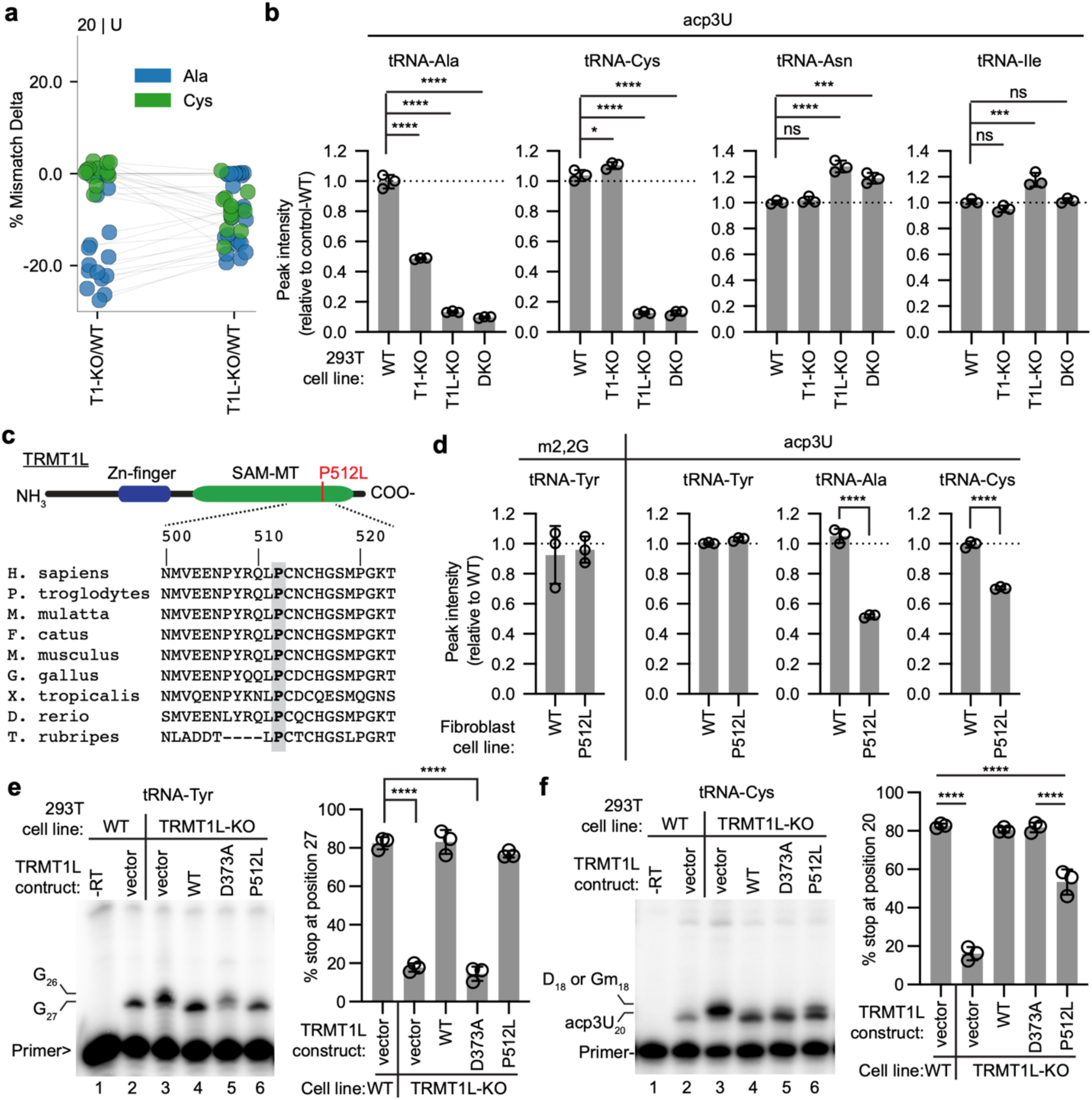
TRMT1 and TRMT1L-deficient human cells exhibit a reduction in acp3U modification in a subset of tRNAs. (a) Change in mismatch incorporation percentage at position 20 for tRNA-Ala or tRNA-Cys isodecoders in the TRMT1-KO or TRMT1L-KO cell line relative to the control-WT cell line. (b) Relative peak intensity values of acp3U modification detected by LC-MS in the indicated tRNAs purified from 293T cell lines versus control-WT. (c) Sequence alignment of vertebrate TRMT1L homologs encompassing the P512 residue. (d) Relative peak intensity values of the indicated modifications detected by LC-MS in purified tRNAs from the TRMT1-KO cell line versus control-WT cell line. (e, f) Representative primer extension assay gel to monitor m2,2G or acp3U in tRNA-Tyr or tRNA-Cys, respectively, from the indicated cell lines with quantification of % stop to the right. Statistical analysis was performed using one-way ANOVA and significance calculated using Tukey’s multiple comparison test. ****P ≤ 0.0001, ***P ≤ 0.001, **P ≤ 0.01.

We next used LC-MS analysis of purified tRNA-Ala and tRNA-Cys to directly measure changes in acp3U modification. We also monitored acp3U in purified tRNA-Asn and tRNA-Ile as comparison. Consistent with the decrease in misincorporation frequency at position 20 of tRNA-Ala detected through OTTR-Seq, we found that the acp3U modification was reduced in tRNA-Ala isoacceptors purified from the TRMT1-KO, TRMT1L-KO, and DKO cell lines (Figure 4b, tRNA-Ala). Also consistent with the tRNA-Seq data, the level of acp3U modification in tRNA-Cys was decreased in the TRMT1L-KO and DKO cell lines, but not the TRMT1-KO cell line (Figure 4b, tRNA-Cys). Interestingly, the TRMT1L-KO cell line exhibited an increase in acp3U modification in tRNA-Asn and tRNA-Ile (Figure 4b, tRNA-Asn and tRNA-Ile, Supplemental Figure 9). We further validated the changes in acp3U modification in the TRMT1-KO, TRMT1L-KO, and DKO cell lines using absolute quantification (Supplemental Figure S10).

To further confirm the changes in acp3U modification, we employed a primer extension assay since acp3U is known to inhibit RT activity (Funk *et al*, 2020; Meyer *et al*, 2020). Since tRNA-Cys does not contain m2,2G at position 26 that would inhibit primer annealing, this allowed us to design a primer hybridizing downstream of position 20 to monitor acp3U status. Using this approach, we detected an RT block at position 20 in tRNA-Cys indicative of the acp3U modification in the control-WT cell line (Supplemental Figure 11). The RT block at position 20 was greatly reduced in the TRMT1L-KO and DKO cell lines, leading to readthrough until the upstream dihydrouridine and/or 2’-O-methylguanosine modification at position 18 (Supplemental Figure 11, lanes 4 and 5). The RT block at position 20 in tRNA-Cys was not greatly affected in the TRMT1-KO cell line (Supplemental Figure 11, lane 3). These results confirm that the acp3U modification at position 20 in tRNA-Cys is affected by loss of TRMT1L expression but not TRMT1. Altogether, these findings suggest that TRMT1 and TRMT1L play a role in modulating acp3U modification at position 20 in the D-loop of tRNAs. Interestingly, TRMT1L impacts the subset of tRNAs whose acp3U modification is dependent upon the DTWD1 enzyme.

### Identification of a pathogenic TRMT1L variant that reduces acp3U but not m2,2G modification in tRNAs

In collaboration with clinicians matched through the GeneMatcher database (Sobreira *et al*, 2015), we identified non-identical sibling patients homozygous for a rare TRMT1L variant who exhibited a range of early-onset neurodegenerative symptoms (Supplemental Figure S12a, patients V.4 and V.5). After initial exome sequencing of the proband proved inconclusive, subsequent analysis revealed a homozygous missense variant in TRMT1L (NM_030934.5): c.1535C>T, p.(Pro512Leu). The variant segregates with the disease in the family and is predicted to be deleterious based upon multiple pathogenicity prediction algorithms (Supplemental Figure S12b and 12c). The P512 residue lies within the methyltransferase domain and is conserved among TRMT1L homologs (Figure 4c).

To investigate the functional impact of this missense variant, fibroblast cells were derived from a skin biopsy taken from one of the siblings (hereafter referred to as P512L patient fibroblast cells). The TRMT1L-P512L patient fibroblast cells were compared to a control fibroblast cell line obtained from a healthy, age-matched individual with wildtype TRMT1L alleles. Using LC-MS, no major change in m2,2G or acp3U levels was detected in purified tRNA-Tyr from the TRMT1L-P512L patient fibroblast cells compared to the control fibroblast cell line (Figure 4d, tRNA-Tyr). In contrast, we detected a decrease in acp3U modification in tRNA-Ala and Cys isolated from the TRMT1L-P512L patient fibroblast cells (Figure 4d). These results suggest that the TRMT1L-P512L variant impairs acp3U modification in tRNA-Ala and Cys without a detectable impact on the m2,2G or acp3U modification in tRNA-Tyr.

We further confirmed and defined the molecular effect of the TRMT1L-P512L variant using our human 293T cell line deficient in TRMT1L. As shown above, we had generated TRMT1L-KO cell lines containing integrated lentiviral vectors expressing either GFP alone, TRMT1L-WT, or the TRMT1L-D373A mutant that exhibits defects in rescuing m2,2G formation in tRNA-Tyr (Figure 2f). Using this same strategy, we generated a TRMT1L-KO cell line stably expressing the TRMT1L-P512L variant (Supplemental Figure S13). As described above, tRNA-Tyr from control-WT cells exhibited a primer extension stop at position 27 while the TRMT1L-KO cell lines with vector alone exhibited background levels of m2,2G stop at position 27 in tRNA-Tyr (Figure 4e, lanes 2 and 3). Re-expression of wildtype TRMT1L, but not TRMT1L-D373A, in TRMT1L-KO cells was able to restore m2,2G formation at position 27 in tRNA-Tyr (Figure 4e, lanes 4 and 5). Consistent with the results in P512L patient fibroblast cells, the re-expression of TRMT1L-P512L in TRMT1L-KO cells was able to restore m2,2G formation at position 27 in tRNA-Tyr (Figure 4e, lane 6).

As shown above, tRNA-Cys from control-WT cells exhibited a RT block at position 20 of tRNA-Cys that was reduced in the TRMT1L-KO cell line with concomitant readthrough to the upstream D18 or Gm18 position (Figure 4f, lanes 2 and 3). Expression of wildtype TRMT1L rescued the RT block at position 20 (Figure 4f, lane 4). Notably, we find that expression of the TRMT1L-D373A mutant is able to rescue acp3U formation in tRNA-Cys despite being impaired in m2,2G formation in tRNA-Tyr (Figure 4f, lane 5). Consistent with the results in P512L patient fibroblast cells, expression of the TRMT1L-P512L variant exhibited only partial rescue of acp3U modification at position 20 in tRNA-Cys of TRMT1L-KO cells (Figure 4f, lane 6). Altogether, these findings uncover a pathogenic variant in TRMT1L that impacts acp3U modification and that TRMT1L function in m2,2G modification can be uncoupled from TRMT1L’s effect on acp3U modification.

### TRMT1-catalyzed modifications impact the levels of tRNA-Tyr and tRNA-Ser in human cells

The above discoveries using OTTR-Seq have revealed a global overview of the modification landscape of tRNAs impacted by TRMT1 and TRMT1L. We next investigated how loss of TRMT1 or TRMT1L affects the abundance of individual tRNAs. We used the AlkB-treated samples for abundance measurements since removing the RT-blocking m^1^A, m^3^C, and m^1^G facilitates more accurate comparison of tRNA levels independent of methyl modification status. As noted above, we observed an increased read coverage for full-length mature tRNA-Tyr-GUA in the TRMT1L-KO cell line caused by increased read-through by the RT due to loss of m2,2G modification at position 27 of tRNA-Tyr (Supplemental Figure S6c). While there was also a slight increase in tRNA-Lys-UUU and tRNA-Phe-GAA in the TRMT1L-KO cell line, no other tRNAs exhibited a significant change in levels in the TRMT1L-KO cell line (Supplemental Figure S6c).

An increase in full-length read abundance was observed for most G26-containing tRNAs in the TRMT1-KO cell line compared to the control-WT cell line (Figure 5a, triangles, Supplemental Figure S14). The increase in read abundance for G26-containing tRNAs in the TRMT1-KO cell line compared to the control-WT cell line is consistent with the lack of m2,2G modification at position 26 in the TRMT1-KO cell line that allows for enhanced RT processivity and more full-length tRNA reads. The apparent increase in full-length tRNA reads in the TRMT1-KO cell line is further amplified by the incomplete removal of the m2,2G modification by AlkB in the WT-control samples that leads to reduced levels of full-length tRNA reads relative to the TRMT1-KO samples.

**Figure 5.**
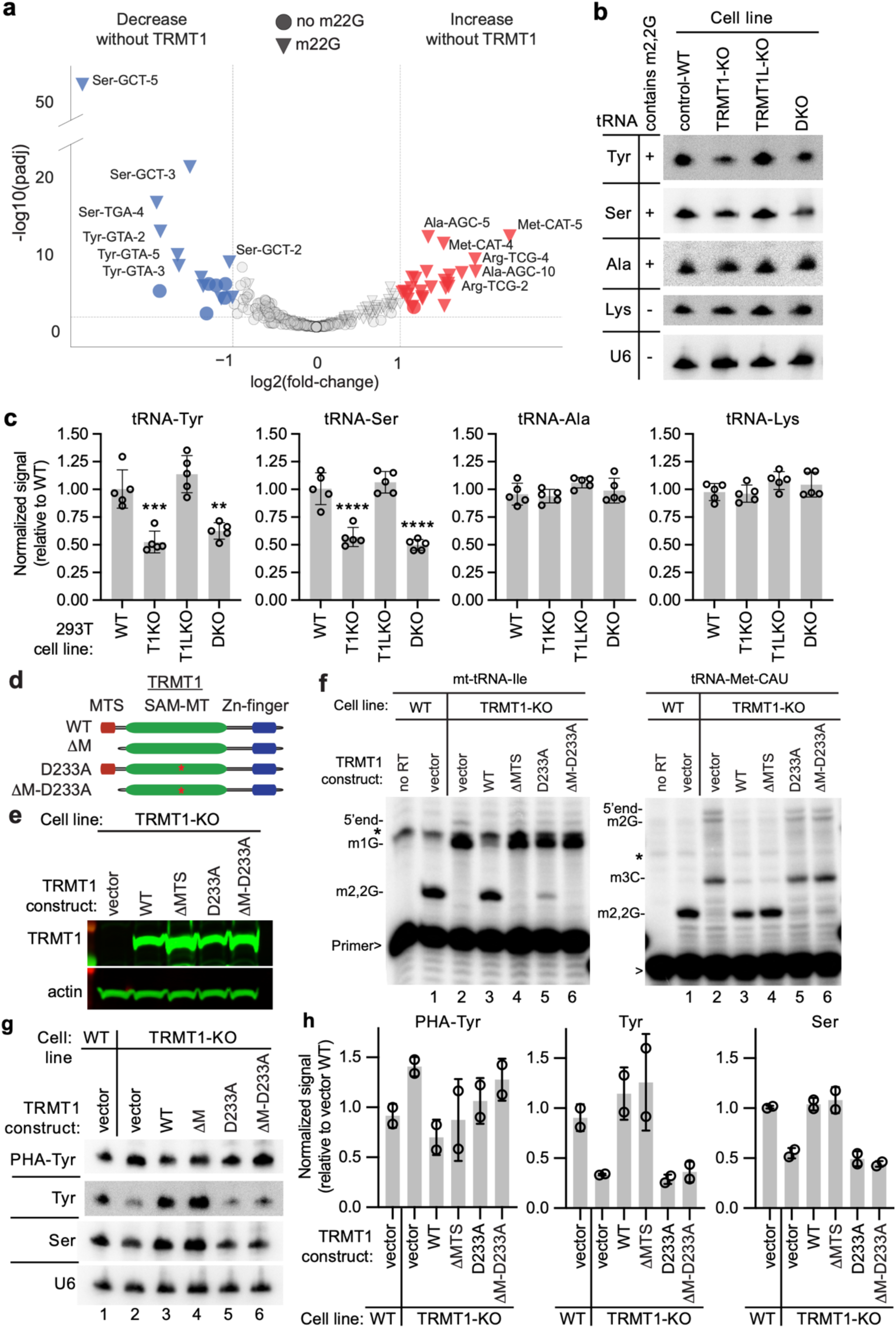
TRMT1-catalyzed modification is required for the accumulation of tyrosine and serine tRNAs in human cells. (a) Volcano plot of tRNA levels in TRMT1-KO versus control-WT cell lines. Triangles represent tRNAs that are modified with m2,2G at position 26. Circles represent tRNAs without m2,2G. Red and blue denote tRNAs that are increased and decreased, respectively, in the TRMT1-KO cell lines. (b) Northern blot analysis of the indicated tRNAs from control-WT, TRMT1-KO, TRMT1L-KO, and TRMT1-TRMT1L-double knockout (DKO) cell lines. (c) Quantification of Northern blots in (b). tRNA levels were normalized using U6 snRNA as a loading control. Bars represent the standard deviation from the mean. Statistical analysis was performed using one-way ANOVA and significance calculated using Dunnett’s multiple comparison test. ****P ≤ 0.0001, ***P ≤ 0.001, **P ≤ 0.01. (d) Schematic of TRMT1-variants used for rescue experiments. (e) Immunoblot analysis confirming expression of TRMT1 variants in the TRMT1-KO cell line. (f) Primer extension analysis of mt-tRNA-Ile and tRNA-Met-CAU from either control-WT or TRMT1-KO cell lines containing stable integration of the indicated TRMT1 constructs. (g) Northern blot analysis of tRNAs from control-WT and TRMT1-KO cell lines containing the indicated TRMT1 construct. (h) Quantification of Northern blots in (g). PHA quantification represents the ratio of PHA versus control probe signal expressed relative to the control-WT cell line.

However, in contrast to most G26-containing tRNAs, we found that all tRNA-Tyr isodecoders and a subset of tRNA-Ser isodecoders exhibited a decrease in abundance in the TRMT1-KO cell line (Figure 5a, blue triangles, Supplemental Figure S14). The decrease in tRNA-Tyr-GUA and tRNA-Ser isodecoders was not observed in TRMT1L-KO cell lines (Supplemental Figure S6c). Both tRNA-Tyr and tRNA-Ser isoacceptors contain m2,2G modification at position 26 that are abolished in the TRMT1-KO cell line. We also measured tRNA modifications in purified tRNA-Ser isoacceptors and detected only background levels of m2,2G in tRNA-Ser purified from TRMT1-KO cells (Supplemental Figure S15). Interestingly, we also found that the m3C, Um and i6A modifications are reduced in tRNA-Ser isolated from TRMT1-KO cells (Supplemental Figure S15). These results suggest a role for TRMT1 in the cellular accumulation of tRNA-Tyr and tRNA-Ser isodecoders as well as maintaining the normal modification patterns present in these tRNAs.

To confirm these findings, we performed Northern blotting using probes against tRNA-Tyr and tRNA-Ser. Consistent with the sequencing data, we find that tRNA-Tyr-GUA isoacceptors exhibited a ∼50 percent decrease in TRMT1-KO cell lines, but not in the TRMT1L-KO cell lines (Figure 5b, c). Further confirming the role of TRMT1 in tRNA-Tyr stability, the TRMT1-TRMT1L-DKO cell lines exhibited a ∼50 percent decrease in tRNA-Tyr similar to the TRMT1-KO cell lines. TRMT1-KO cells also exhibited a ∼50 percent decrease in tRNA-Ser-GCU while TRMT1L-KO cell lines displayed similar levels of tRNA-Ser-GCU as control-WT cells (Figure 5b, c). The DKO cell line exhibited a similar reduction in tRNA-Ser-GCU as TRMT1-KO cell lines providing evidence that TRMT1-deficiency contributes to nearly all the reduction of tRNA-Ser, like tRNA-Tyr. We also probed against tRNA-Ala isoacceptors, which is another m2,2G-containing tRNA, but did not detect a significant change in levels between any of the cell lines (Figure 5b, c). We also probed against tRNA-Lys-UUU, which does not contain m2,2G, and observed no major change in abundance (Figure 5b, c).

We next investigated whether the presence of TRMT1 protein alone and/or the m2,2G modification are required for tRNA stability. To test whether the activity of TRMT1 is required for tRNA stability, we performed rescue experiments using re-expression of wildtype TRMT1 or a TRMT1 variant containing an D233A mutation in the SAM binding domain of the methyltransferase domain that is expected to reduce catalytic activity (Figure 5d). We also created a construct expressing a TRMT1 variant lacking the N-terminal mitochondrial targeting signal without or with the D233A mutation (Figure 6a, ΔM, ΔM-D233A). We generated stable cell lines in the TRMT1-KO background and confirmed re-expression of TRMT1 by immunoblotting (Figure 5e).

**Figure 6.**
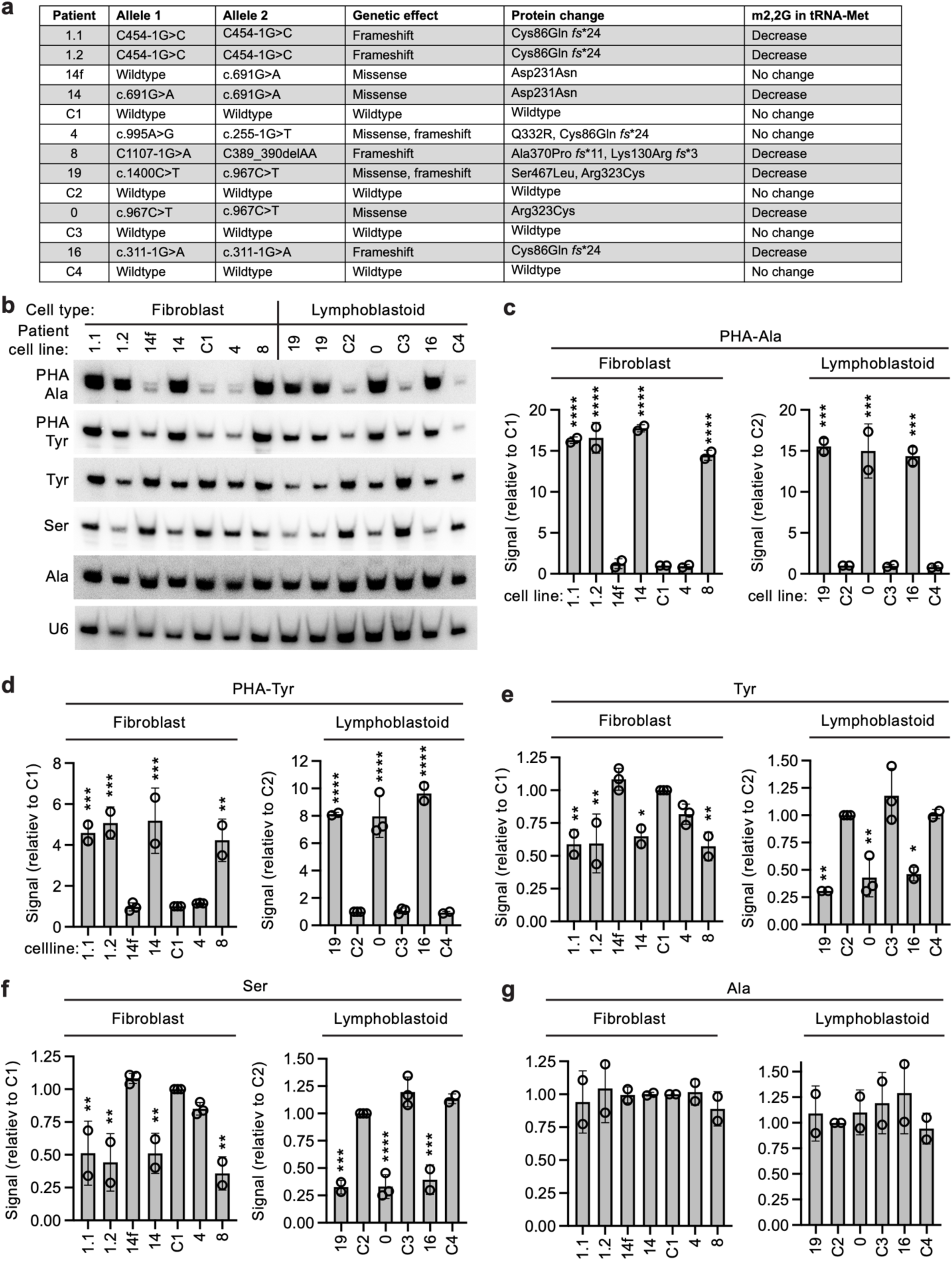
Human patient cells containing biallelic pathogenic TRMT1 variants exhibit a reduction in serine and tyrosine tRNA levels. (a) Patient and control cell lines used for analysis along with genotypes, expected genetic outcome, and known effects on TRMT1 protein and m2,2G levels. (b) Northern blot analysis of fibroblast or lymphoblastoid cell lines derived from patient and control individuals. (c, d, e, f, and g) Quantification of the relative Northern blot signal for the indicated tRNAs from patient cell line versus the C1 control fibroblast cell line or C2 control lymphoblastoid cell line. PHA quantification represents the ratio of PHA versus control probe signal expressed relative to either the C1 or C2 control cell line. Error bars indicate the standard deviation from the mean. Statistical analysis was performed using one-way ANOVA and significance calculated using Dunnett’s multiple comparison test. ****P ≤ 0.0001, ***P ≤ 0.001, **P ≤ 0.01, *P ≤ 0.05.

To validate the functional re-expression of TRMT1, we monitored the rescue of m2,2G formation in tRNA-Met-CAU and mt-tRNA-Ile-GAU. As expected, TRMT1-KO cells with vector alone exhibited no m2,2G stop at position 26 in tRNA-Met-CAU or mt-Ile-GAU when compared to wildtype control cells (Figure 5f, compare lanes 1 and 2). Re-expression of wildtype TRMT1 in TRMT1-KO cells was able to rescue m2,2G formation in tRNA-Met-CAU or mt-Ile-GAU as evidenced by the re-introduction of the RT block at position 26 and the reduction of read-through products (Figure 5f, lane 3). The re-expression of TRMT1-ΔM was able to restore m2,2G modification in tRNA-Met-CAU, but not mt-tRNA-Ile-GAU (Figure 5f, lane 4). Expression of TRMT1-D233A or ΔM-D233A in TRMT1-KO cell lines led to only minor restoration of m2,2G formation at position 26 in either tRNA-Met-CAU or mt-tRNA-Ile-GAU (Fig. 6c, lanes 5 and 6).

Due to the m2,2G modification catalyzed by TRMT1L at position 27 in tRNA-Tyr, the m2,2G status at the upstream position 26 cannot be ascertained by primer extension in TRMT1-KO cell lines. Instead, we employed the PHA assay described above to monitor m2,2G formation at position 26. Using the PHA assay, we detected an increase in PHA probe signal for tRNA-Tyr-GUA in the TRMT1-KO cell line compared to control-WT cells (Figure 5g, compare lanes 1 and 2, quantified in 5h). Re-expression of TRMT1 or TRMT1-ΔM in the TRMT1-KO cell lines resulted in a reduced PHA signal indicative of m2,2G restoration compared to TRMT1-KO cells with vector (Figure 5g, compare lane 2 to lanes 3 and 4, quantified in 5h). In contrast, expression of the TRMT1-D233A or TRMT1-ΔM-D233A variant led to only partial reduction in PHA signal in tRNA-Tyr (Figure 5g, lanes 5 and 6, quantified in 5h). These results further validate that TRMT1 catalyzes the formation of m2,2G at position 26 in tyrosine tRNAs.

We next monitored the accumulation of mature tRNA-Tyr and Ser in the TRMT1-KO cell lines re-expressing TRMT1 variants. We find that TRMT1-KO cells with vector alone exhibited decreased levels of tRNA-Tyr and tRNA-Ser relative to the control-WT strain (Fig. 5g, tRNA-Tyr row, compare lanes 1 and 2, quantified in 5h). Re-expression of wildtype TRMT1 or TRMT1-ΔMTS in TRMT1-KO cells was able to rescue tRNA-Tyr and tRNA-Ser levels back to wildtype control levels (Fig. 5g, compare lane 2 to lanes 3 and 4, quantified in 5h). In contrast, expression of the TRMT1 variants with mutations in the active site were unable to rescue tRNA-Tyr or tRNA-Ser levels and remained at similar levels as the TRMT1-KO cell lines with empty vector (Fig. 5g, lanes 5 and 6, quantified in 5h). Altogether, these results uncover a key role for TRMT1-catalyzed m2,2G modifications in the cellular stability of tRNA-Tyr and Ser isoacceptors.

### Human cells with pathogenic TRMT1 variants exhibit perturbations in tRNA-Tyr and Ser levels

Homozygosity of pathogenic variants in *TRMT1* have been shown to cause nearly complete loss of m2,2G modifications in human patients with intellectual disability disorders (Jonkhout *et al*., 2021; Zhang *et al*., 2020). Our findings described above indicate that a decrease in m2,2G modifications could impact the levels of tRNA-Tyr and tRNA-Ser in human patient cells within a disease setting. Thus, we monitored the levels of tRNA-Tyr and tRNA-Ser in human patient cell lines containing biallelic TRMT1 variants that are associated with intellectual disability disorder (Figure 6a) (Blaesius *et al*., 2018; Davarniya *et al*., 2015; Jonkhout *et al*., 2021; Monies *et al*., 2017; Najmabadi *et al*., 2011; Zhang *et al*., 2020) (Efthymiou et al., *manuscript in preparation*). Patients 1.1, 1.2, 14, 8, 19, 0, and 16 in this panel have inherited TRMT1 variants that cause aberrant splicing, reduced TRMT1 expression, and/or loss of TRMT1 activity (Figure 6a, shaded). Patient 4 is heterozygous for a non-pathogenic TRMT1 missense allele that does not impact m2,2G levels while Patient 14f is the heterozygous father of patient 14 who contains a wildtype allele of TRMT1 and does not exhibit a major change in m2,2G modification levels. The patient cell lines were compared to control cell lines from four healthy, unrelated individuals containing wildtype alleles of TRMT1 (Figure 6a, C1 to C4).

We first validated that m2,2G levels were decreased in the patient cell lines using the PHA assay against tRNA-Ala. All patient cell lines except for the patient 14f and 4 cell lines exhibited an increased PHA signal for tRNA-Ala relative to the control patient samples (Figure 6b, quantified in 6c). In addition, we found that all patient cell lines except for patient 4 and 14f exhibited an increase in PHA signal for tRNA-Tyr relative to control cell lines indicative of loss of m2,2G formation at position 26 in tRNA-Tyr (Figure 6b and 6d). Notably, we found that tRNA-Tyr and tRNA-Ser levels are reduced in all patient cell lines exhibiting deficits in m2,2G modification caused by pathogenic TRMT1 variants (Figure 6b, quantified in 6e and 6f). Consistent with our findings above that TRMT1 does not affect tRNA-Ala levels, no significant changes in tRNA-Ala were detected between any of the patient versus control cell lines (Figure 6b and 6g). These findings show that loss of m2,2G modification across a spectrum of individuals with pathogenic TRMT1 variants all impact serine and tyrosine tRNA levels in a physiological context of human neurodevelopmental disorders.

### TRMT1 and TRMT1L are required for efficient translation of tyrosine and serine codons

The preceding results suggest that TRMT1- and/or TRMT1L-catalyzed tRNA modifications could affect the translation of tyrosine and serine codons by ensuring proper levels of functional tRNA-Tyr and tRNA-Ser. Thus, we investigated the possible effects of TRMT1 and TRMT1L on protein synthesis using codon-dependent reporter constructs. We generated mammalian expression reporters encoding nanoluciferase protein fused downstream of ten consecutive tyrosine or serine codons (Figure 7a). The reporter constructs also contain a cassette encoding firefly luciferase expressed from a separate promoter to control for transfection efficiency (Figure 7a). The nanoluciferase signal from the codon-dependent reporter was then normalized against an additional reporter without the serine or tyrosine codon stretch to control for any differences in translation independent of the tyrosine or serine codons. Expression was confirmed by immunoblotting (Supplemental Figure S16).

**Figure 7.**
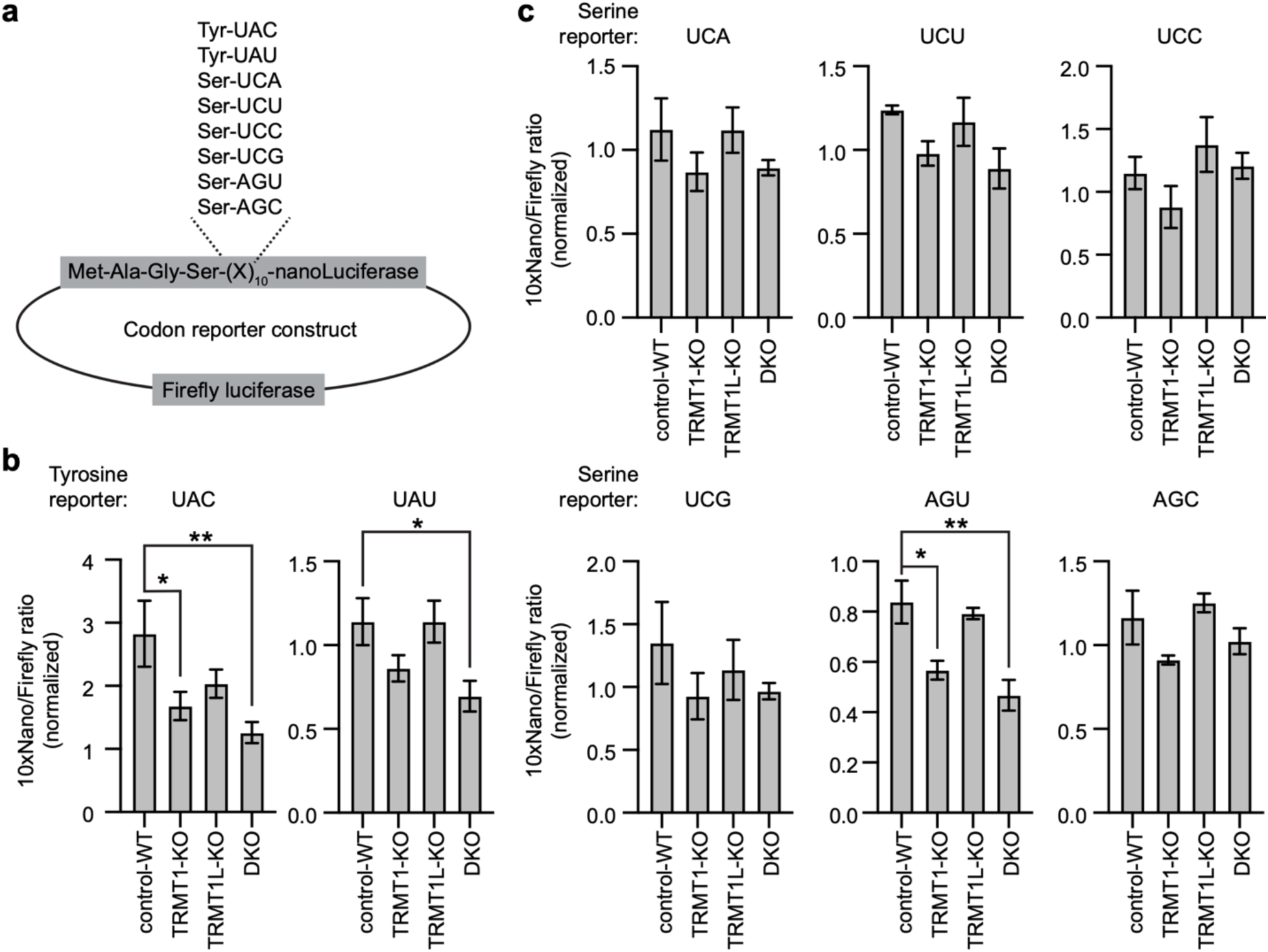
Loss of TRMT1 and TRMT1L impairs protein expression from specific codon reporters. (a) Schematic of plasmid reporter constructs. Plasmids encode nanoLuciferase fused in frame with ten consecutive tyrosine or serine codons. Firefly luciferase is expressed from the same plasmid to serve as an internal control for transfection efficiency. (b) Quantification of protein expression from the tyrosine UAC and UAU codon reporters. (c) Quantification of translation from the indicated serine codon reporters. For (b) and (c), the signal from the nanoluciferase reporter protein (Nano) is expressed relative to firefly luciferase protein following normalization to a control reporter without the serine or tyrosine codon stretch. Error bars indicate the standard error from the mean. Statistical analysis was performed using one-way ANOVA and significance calculated using Dunnett’s multiple comparison test. **P ≤ 0.01, *P ≤ 0.05.

Based upon this system, we found that TRMT1-KO cells exhibited a decrease in expression of the tyrosine UAC and UAU codon reporter relative to control-WT cells with the UAC reporter exhibiting a significant reduction (Figure 7b, TRMT1-KO). We also observed a decrease in translation of the UAC tyrosine codon reporter in the TRMT1L-KO cell line, but this difference did not reach statistical significance (Figure 7b, TRMT1L-KO). The TRMT1-TRMT1L DKO cell line exhibited an even further reduction in translation of the tyrosine UAC and UAU codon reporters, indicative of an additive decrease when both TRMT1 and TRMT1L are absent (Figure 7b, DKO).

Similar to the tyrosine codon reporters, the TRMT1-KO cell line exhibited a reduction in translation for each of the serine codon reporters that was most pronounced with the Serine AGU codon reporter (Figure 7c, TRMT1-KO). We note that the AGU codon is decoded by the tRNA-Ser-GCU isoacceptor family, which exhibits decreased levels in the TRMT1-KO cell line. The TRMT1L-KO cell line displayed a similar level of expression for each of the serine codon reporters as the control-WT cell line (Figure 7c, TRMT1L-KO). The lack of change is consistent with a role for TRMT1L in the modification of tRNA-Tyr, but not tRNA-Ser isoacceptors. Except for the serine UCC reporter, the DKO cell lines exhibited a similar decrease in translation for the serine codon reporters as the TRMT1-KO cell line (Figure 7c, DKO). Altogether, these results suggest that the decrease in tRNA-Tyr and tRNA-Ser levels in TRMT1-KO cells reduces translation efficiency of cognate codons while TRMT1L-catalyzed modification has an additive effect with TRMT1 on the activity of tRNA-Tyr isoacceptors.

## Discussion

While the presence of m2,2G at positions 26 and 27 in mammalian tyrosine tRNAs has been known for decades, the enzyme responsible for modifying guanosine at position 27 has been elusive (Johnson *et al*., 1985; Liu & Straby, 2000; van Tol *et al*., 1987). Here, we demonstrate that TRMT1L catalyzes m2,2G formation at position 27 in human tyrosine tRNAs. Importantly, and concurrent with the work described here, Hwang *et al*. also report that human TRMT1L is required for m2,2G modification in tyrosine tRNAs and plays an unexpected role in acp3U modification (Hwang *et al*, 2024). Together, our parallel studies share and corroborate each other’s main findings on TRMT1L and its substrates.

TRMT1L orthologs have been identified in all sequenced vertebrates, but not in invertebrates, plants, or single-cell organisms (Jonkhout *et al*., 2021; Towns & Begley, 2012). TRMT1L could have arisen from a gene duplication event in vertebrates from an ancestral Trm1 homolog and evolved to modify position 27 in tyrosine tRNAs. Consistent with this hypothesis, tyrosine tRNAs in insects and plants contain G at position 27 but are not modified with m2,2G (Cappannini *et al*., 2023; Funk *et al*., 2020). Interestingly, TRMT1L-deficient mice exhibit altered motor coordination and aberrant behavior without any major anatomical changes (Vauti *et al*, 2007). Moreover, TRMT1L represents one of several genes that has undergone shifts in expression within the cerebellum of certain mammals (Brawand *et al*, 2011). TRMT1 and TRMT1L also exhibit changes in their subcellular localization in human neural cells upon neuronal activation (Jonkhout *et al*., 2021). These findings suggest that the advent of TRMT1L and changes in its expression pattern could underlie alterations in motor control function among vertebrates. Our identification of a pathogenic TRMT1L variant associated with motor neuropathy and leukodystrophy in human patients that perturbs tRNA modification provides key support for this conclusion and uncovers a role for acp3U modification in the nervous system.

Based upon results presented here and the localization of TRMT1 in the nucleus of human cells (Dewe *et al*., 2017), we propose that TRMT1 catalyzes m2,2G formation at position 26 as one of the earliest modifications in unspliced pre-tRNA-Tyr isoacceptors. The presence of m2,2G at position 26 in pre-tRNA-Tyr is consistent with studies showing that pre-tRNA-Tyrosine with an intron contains m2,2G (Laski *et al*, 1983; Nishikura & De Robertis, 1981) and that TRMT1, but not TRMT1L, can modify intron-containing pre-tRNA-Tyr as shown here. The TRMT1-catalyzed modification in pre-tRNA-Tyr could prevent misfolding and stabilize the unspliced tRNA against degradation. After modification of pre-tRNA-Tyr by TRMT1 and intron splicing, then TRMT1L can modify G27 in the spliced tRNA-Tyr. The m2,2G at position 27 could further refine the folding of tRNA-Tyr, thereby promoting the formation of additional modifications and adoption of a functional conformation.

While more than half of all tRNA isodecoders are modified by TRMT1, our studies reveal tyrosine and serine tRNAs as being exquisitely dependent upon TRMT1-catalyzed m2,2G modification for stability and accumulation. This result mirrors findings in yeast showing that destabilization of the acceptor and T-stems of tRNA-Ser or tRNA-Tyr cause recognition and degradation by the rapid tRNA decay (RTD) pathway (Guy *et al*, 2014; Whipple *et al*, 2011). The presence of m2,2G modification at position 26 or 27 of tRNA-Tyr could be important for preventing alternative folding patterns as proposed previously (Steinberg & Cedergren, 1995). In the absence of TRMT1-catalyzed tRNA modifications, tyrosine tRNAs might fold into alternative conformations with destabilized stems that are refractory to further modifications, leading to the recognition and degradation of the hypomodified tyrosine tRNAs. In the case of tRNA-Ser, loss of TRMT1-dependent m2,2G correlates with a reduction in m3C, Um, i6A, and m2G. In yeast, loss of Trm1 also leads to a decrease in m3C modification at position 32 in tRNA-Ser-UGA as well as an increase in m1G at position 9 in a subset of tRNAs (Behrens *et al*., 2021; Hernandez-Alias *et al*., 2023). In both yeast and humans, the m2,2G modification could influence the orientation of the anticodon loop that contains both the i6A and m3C modifications to ensure efficient recognition by their respective tRNA modification enzymes. It will be interesting to determine the pathways that recognize hypomodified serine and tyrosine tRNAs in human cells, since much less is known about tRNA surveillance pathways outside of yeast and bacteria.

Notably, the levels of acp3U modification in certain tRNAs appears to be sensitive to changes in TRMT1 or TRMT1L. In the case of TRMT1, the m2,2G modification catalyzed by TRMT1 could influence the folding and orientation of the D-loop that affects the addition of acp3U at position 20 by the DTWD1/2 enzymes (Takakura *et al*., 2019). We note prior studies in *E. coli* which find that TrmB catalyzed methylation of G46 to m7G promotes acp3U formation at position U47 (Meyer *et al*., 2020). TRMT1L could also facilitate acp3U formation in tRNA-Ala and tRNA-Cys through interaction with the tRNA substrates of DTWD1 and/or modulation of DTWD1 activity since neither tRNA-Ala nor tRNA-Cys contain G at position 27 for modification by TRMT1L. Our studies suggest the intriguing possibility that TRMT1L might exert separable activities required for m2,2G and acp3U modification in distinct tRNA substrates. Alternatively, TRMT1L could play an indirect role through changes in gene expression that reduce DTWD1 protein levels or factors that regulate DTWD1 activity. Future studies will investigate this unexpected expansion of functions for TRMT1L.

Our findings using reporter assays indicate that TRMT1 and TRMT1L-catalyzed modifications in tRNA can influence the efficient translation of certain tyrosine and serine codons. The perturbations in tyrosine and serine tRNA levels could impact protein synthesis and contribute to the reduced levels of global translation detected in TRMT1-KO and TRMT1L-KO cell lines (Dewe *et al*., 2017; Hwang *et al*., 2024). Interestingly, depletion of tyrosine tRNAs results in the impaired translation of growth and metabolic genes enriched in cognate tyrosine codons leading to repressed proliferation (Huh *et al*, 2021). The repressed proliferation associated with tRNA-Tyr depletion is consistent with the growth defect that we have detected in TRMT1-knockout human cell lines as well as the developmental delays observed in human patients with TRMT1- or TRMT1L-deficiency (Blaesius *et al*., 2018; Davarniya *et al*., 2015; Dewe *et al*., 2017; Monies *et al*., 2017; Najmabadi *et al*., 2011). By pinpointing specific tRNAs and their mechanism of dysregulation, the results presented here could guide tRNA-based strategies for rescuing phenotypes associated with neurodevelopmental disorders linked to tRNA modification deficiency (Anastassiadis & Kohrer, 2023; Hou *et al*, 2024).

## Limitations of study

The OTTR-Seq method can only detect a subset of modifications that cause miscoding. Thus, there could be additional modifications impacted by TRMT1 and/or TRMT1L that are missed in this study. Future studies based upon advances in Nanopore sequencing could address this limitation. While we confirmed that tRNA-Tyr and Ser levels are decreased in TRMT1-ID patient cells, the impact of TRMT1 and TRMT1L on additional tRNAs in other tissues and developmental stages is unknown. Finally, the study does not address the potential effects of TRMT1- and TRMT1L-modifications on the translation of codons besides tyrosine and serine codons. The use of ribosome profiling and proteomics will be the subject of future investigation to examine the global effects of TRMT1 and TRMT1L on gene expression.

## Supplemental information

Document S1. Materials and Methods. Supplemental Figures S1–S16, Tables S1-S4, and Supplemental Data 1.

## Acknowledgements

We thank Kevin Welle and the University of Rochester Mass Spectrometry Resource Laboratory for assistance in mass spectrometry. We thank Dr. Eric Phizicky and members of the Fu Lab for feedback on this manuscript. This work was funded by NSF CAREER Award 1552126 and NIH GR530882 to D.F; the Deutsche Forschungsgemeinschaft [325871075-SFB 1309 and 259130777-SFB 1177] to S.M.K.; the Wellcome Trust (WT093205MA, WT104033AIA), the Medical Research Council (MR/S01165X/1, MR/S005021/1, G0601943), The National Institute for Health Research University College London Hospitals Biomedical Research Centre, Rosetrees Trust, Ataxia UK, Multiple System Atrophy Trust, Brain Research United Kingdom, Sparks Great Ormond Street Hospital Charity, Muscular Dystrophy United Kingdom (MDUK), Muscular Dystrophy Association (MDA USA) to H.H.. In addition, S.E. and H.H. were supported by an MRC strategic award to establish an International Centre for Genomic Medicine in Neuromuscular Diseases (ICGNMD) MR/S005021/1.

## Author contributions

Conceptualization, K.Z, J.M.L,, and D.F.; Software, Formal Analysis, Data Curation, and Visualization, A.C.M. and P.C.; Investigation, K.Z., J.M.L., R.M., F.D., J.H., S.I.E., G.Y., H.C., and E.G.K.; Resources, B.B., S.E.; Writing – Original Draft, D.F.; Writing – Review & Editing, K.Z., A.M., R.M., F.D., S.M.K., T.M.L., and D.F.; Funding Acquisition and Supervision, H.H., S.M.K., T.M.L., and D.F.

