## Supplemental Information for "Human TRMT1 and TRMT1L paralogs ensure the proper modification state, stability, and function of tRNAs"

**Zhang et al. 2024**

### Materials and Methods

#### Tissue cell culture

The 293T human embryonic cell line was obtained from ATCC (CRL-3216). Human lymphoblastoid cell lines were generated by EBV immortalization of primary human lymphocytes obtained from patient blood samples. Human fibroblast cell lines were obtained from patients by skin biopsy. 293T human embryonic kidney cell lines and fibroblast cell lines were cultured in Dulbecco's Minimal Essential Medium (DMEM) supplemented with 10% fetal bovine serum (FBS), 1X penicillin and streptomycin (ThermoFisher), and 1X Glutamax (Gibco) at 37 °C with 5% $CO_2$ and ambient 20% $O_2$. Patient fibroblast cells were maintained at 5% $O_2$. Cells were passaged every 3 days with 0.25% Trypsin. Human lymphoblastoid cell lines (LCL) were cultured in Roswell Park Memorial Institute (RPMI) 1640 Medium containing 15% fetal bovine serum, 2 mM L-alanyl-L-glutamine (GlutaMax, Gibco) and 1% Penicillin/Streptomycin at 37 °C with 5% $CO_2$ and 5% $O_2$. LCLs were passaged every 3 days by dilution.

#### Generation of CRISPR-edited Cell Lines

CRISPR editing was performed using plasmid pX333 (Andrea Ventura, Addgene plasmid # 64073) [1]. The oligonucleotide DNA sequences encoding single guide RNA (sgRNA) targeting TRMT1 and TRMT1L were cloned into px333 using BsmB1 and BbsI (Supplemental Table 3). Plasmids were transfected into human 293T embryonic cells using calcium phosphate transfection. Individual clones were by single-cell depositing into 96-well plates. The presence of CRISPR-induced deletion in either TRMT1 or TRMT1L was detected by PCR amplification and confirmed by Sanger sequencing of PCR products amplified from genomic DNA using primers TRMT1 gPCR F2 and TRMT1 gPCR R6 (Supplemental Table 3).

**Immunoblot assays**

Cell extracts and protein samples were loaded onto BOLT 4–12% Bis-Tris gels (ThermoFisher) followed by immunoblotting onto Immobilon-FL PVDF membrane (Millipore Sigma IPFL00010). Antibodies are listed in Supplemental Table 3. Image analysis of immunoblots were performed using Image Studio software (Li-Cor).

**OTTR-seq Library Preparation**

RNA 3' de-phosphorylation was carried out as previously described [2] using 500 ng of total RNA from each sample. Briefly, samples were treated with T4 Polynucleotide Kinase (T4 PNK; New England Biolabs) in a modified 5x reaction buffer (350 mM Tris-HCl, pH 6.5, 50 mM MgCl2, 5 mM dithiothreitol) under low pH conditions in the absence of ATP for 30 minutes. OTTR-seq libraries were generated as previously described [3]. Briefly, total PNK-treated RNA was 3' tailed using mutant BoMoC RT in buffer containing only ddATP for 90 minutes at 30℃, with the addition of ddGTP for another 30 minutes at 30℃. This was then heat-inactivated at 65℃ for 5 minutes, and unincorporated ddATP/ddGTP was hydrolyzed by incubation in 5 mM MgCl2 and 0.5 units of shrimp alkaline phosphatase (rSAP) at 37℃ for 15 minutes. 5 mM EGTA was added and incubated at 65℃ for 5 minutes to stop this reaction. Reverse transcription was then performed at 37℃ for 30 minutes, followed by heat inactivation at 70℃ for 5 minutes. The remaining RNA and RNA/DNA hybrids were then degraded using 1 unit of RNase A at 50℃ for 10 minutes. cDNA was then cleaned up using a MinElute Reaction CleanUp Kit (Qiagen). To reduce adaptor dimers, cDNA was run on a 9% UREA page gel, and the size range of interest was cut out and eluted into gel extraction buffer (300 mM NaCl, 10 mM Tris; pH 8.0, 1 mM EDTA, 0.25% SDS) and concentrated using EtOH precipitation. Size-selected cDNA was then PCR amplified for 12 cycles using Q5 High-fidelity polymerase (NEB #M0491S). Amplified libraries were then run on a 6% TBE gel, and the size range of interest was extracted to reduce adaptor dimers further. Gel slices were eluted into gel extraction buffer (300 mM NaCl, 10 mM Tris; pH 8.0, 1 mM EDTA) followed by concentration using

EtOH precipitation. Final libraries were pooled and sequenced on an Illumina NextSeq 500 150-cycle high-output kit.

**Data Processing**

Sequencing adaptors were trimmed from raw reads using cutadapt, v1.18, and read counts were generated for small RNA types using tRAX [4]. Briefly, trimmed reads were mapped to the mature tRNAs of GRCh38/hg38 human genome assembly obtained from GtRNAdb [5,6] and the reference genome sequence using Bowtie2 in very-sensitive mode with the following parameters to allow for a maximum of 100 alignments per read: --very-sensitive --ignore-quals --np 5 -k 100. Mapped reads were filtered to retain only the "best mapping" alignments. Raw read counts of tRNAs and other small RNA types were computed using tRNA annotations from GtRNAdb, and annotations from GENCODE M23 and miRBase v22 [7]. Raw read counts were then normalized using DESeq2. Downstream data visualizations were generated with custom Python scripts using the standard tRAX output files as input. Primary data can be accessed on the GEO database.

**Modification-induced misincorporations**

To determine sites of potential tRNA modifications present in the sequencing data, a custom Python script was used with the tRAX output files as the input (https://github.com/Aimann/tMAP). Briefly, tRNA transcripts were filtered to have at least 20 reads encompassing the length of the tRNA transcript and must contain >=80% uniquely mapped reads to an individual tRNA. Per-base read coverage was calculated for each of these transcripts and the counts of adenines, guanines, cytosines, thymines, and deletions were summed, and the number of non-reference bases was calculated. The number of non-reference bases was then divided by the total read coverage at that position to determine the misincorporation rate at that position. Downstream data visualizations were generated with a custom python script using these mismatch values for each tRNA, across each tissue.

**RNA analysis**

RNA was extracted using TRIzol LS reagent (Invitrogen). RNAs were diluted into formamide load buffer, heated to 95°C for 3 minutes, and fractionated on a 10% polyacrylamide, Tris-Borate-EDTA (TBE) gel containing 7 M urea. Sybr Gold nucleic acid staining (Invitrogen) was conducted to identify the RNA pattern. RNA was subsequently transferred to a Hybond N+ membrane (GE Healthcare Life Sciences) for probe hybridization with radiolabeled oligonucleotides (Supplemental Table 3). Blots were stripped by incubation at 75°C with stripping buffer (0.15 M NaCl, 0.015 M Na-citrate, 0.1% SDS) and repeated at least twice until there was no detectable signal.

For primer extension analysis, 1.5 µg of total RNA was pre-annealed with 5'-$^{32}$P-labeled oligonucleotide and 5x hybridization buffer (250 mM Tris, pH 8.5, and 300 mM NaCl) in a total volume of 7 µl. The mixture was heated at 95°C for 3 min followed by slow cooling to 42°C. An equal amount of extension mix consisting of avian myeloblastosis virus reverse transcriptase (Promega), 5x AMV buffer and 40 µM dNTPs was added. The mixture was then incubated at 42°C for 1 hour and loaded on 15% 7 M urea denaturing polyacrylamide gel. Gels were exposed on a phosphor screen (GE Healthcare) and scanned on a Bio-Rad personal molecular imager or Azure Sapphire Biomolecular Imager followed by analysis using NIH ImageJ software. Primer extension oligonucleotide sequences were previously described [8] or in Supplemental Table 3.

**Purification of tRNAs**

RNA was extracted from HEK 293T cell lines by TRIzol LS reagent. Small RNA (<200 nucleotides) was then purified using RNA Clean and Concentrator Zymo-Spin IC Columns (Zymo Research). RNA and 5' biotin-oligos (Supplementary Table 3) were boiled in of 95 °C for 3 min in hybridization buffer (60mM NaCl, 50mM Tris pH8.5), followed by slow cooling to 25 °C (room temperature). Dynabeads MyOne Streptavidin C1 (Invitrogen, 65001) was washed in 6X SSC (0.9M Sodium chloride, 90 mM sodium citrate), then applied to the RNA-oligo mixture for 3 hours at 25 °C (room temperature). Beads were separated from the flowthrough using a magnetic rack, and the beads were washed 3 times in 3xSSC followed by 2 times in 1xSSC. Then RNase-free water was added to elute the purified RNA from the resin at 65 °C. The elution step was repeated 3 times and combined for LC-MS analysis.

**Liquid chromatography-mass spectrometry**

RNAs were digested and processed by LC-MS as described previously [9]. Briefly, ribonucleosides were separated using a Hypersil GOLDTM C18 Selectivity Column (Thermo Scientific) followed by nucleoside analysis using a Q Exactive Plus Hybrid Quadrupole-Orbitrap. The modification ratio was calculated using the m/z intensity values of each modified nucleoside following normalization to the sum of intensity values for the canonical nucleosides; A, U, G and C.

For molar quantification of RNA modifications in Supplemental Figure X, we enzymatically digested isolated isoacceptors to their nucleoside form using a mixture of Sigma Aldrich Bioultra Alkaline Phosphatase from bovine intestinal mucosa, Sigma Aldrich Benzonase Nuclease and Merck Phosphodiesterase I. Digestion was performed for 2h at 37°C and digests were filtered through a 10 kDa MWCO plate filter at 4°C and 3000 rcf for 15 minutes. Samples were coinjected with 1 µL of stable isotope labelled internal standard (SILIS) derived from metabolically labelled yeast tRNA as described previously [10]. Modified nucleosides were identified and quantified using mass spectrometry. An Agilent 1290 Infinity II equipped with a diode-array detector (DAD) combined with an Agilent Technologies G6470A Triple Quad system and electrospray ionization (ESI-MS, Agilent Jetstream) was used.

Nucleosides were separated using a Synergi Fusion-RP column (Synergi® 2.5 µm Fusion-RP 100 Å, 150 × 2.0 mm, Phenomenex®, Torrance, CA, USA). LC buffer consist of 5 mM $NH_4OAc$ pH 5.3 (buffer A) and pure acetonitrile (buffer B) were used as buffers. The gradient starts with 100% buffer A for 1 min, followed by an increase to 10% buffer B over a period of 4 min. Buffer B is then increased to 40% over 2 min and maintained for 1 min before switching back to 100% buffer A over a period of 0.5 min and re-equilibrating the column for 2.5 min. The total time is 11 min and the flow rate is 0.35 mL/min at a column temperature of 35 °C.

An ESI source was used for ionization of the nucleosides (ESI-MS, Agilent Jetstream). The gas temperature (N2) was 230 °C with a flow rate of 6 L/min. Sheath gas temperature was 400 °C with a flow rate of 12 L/min. Capillary voltage was 2500 V, skimmer voltage was 15 V, nozzle voltage was 0 V, and nebulizer pressure was 40 Psi. The cell accelerator voltage was 5 V. For quantification, a DMRM and positive ion mode was used (Supplemental Table S4).

**Plasmids**

The pcDNA3.1-TRMT1-FLAG construct was previously described [8]. The open reading frame for TRMT1 was cloned into pcDNA3.1-Twin-Strep to yield pcDNA3.1-TRMT1-Strep. The ORF for TRMT1L (NM_030934.5) was PCR amplified from cDNA plasmid HsCD00337317 (Plasmid Repository, Harvard Medical School) and cloned into either pcDNA3.1-TWIN-Strep or pcDNA3.1-3xFLAG. The pcDNA3.1-TRMT1-Strep-D233A and pcDNA3.1-Strep-TRMT1L-D373A expression constructs were generated by DpnI site-directed mutagenesis using oligonucleotides in Supplemental Table 3. All plasmid constructs have been verified by whole plasmid sequencing (Plasmidsaurus).

For lentiviral expression, the open reading frame for TRMT1 or TRMT1-D233A was cloned into pLenti CMV GFP Blast (659-1)[11]. For lentiviral expression of TRMT1L, the open reading frame for TRMT1L or TRMT1L-D373A was PCR amplified and cloned into pLenti CMV GFP Puro (658-5). pLenti CMV GFP Blast (659-1) (Addgene plasmid #17445) and pLenti CMV GFP Puro (658-5) (Addgene plasmid #17448) were gifts from Eric Campeau and Paul Kaufman. Cloning was performed using the T5 exonuclease-dependent assembly method [12].

**Generation of stable cell lines**

To generate lentivirus, $2.5 \times 10^5$ 293T cells were seeded onto 60 × 15 mm tissue culture dishes. Then, 1.25 µg of lentiviral transfer plasmids along with a lentiviral packaging cocktail containing 0.75 µg of psPAX2 packaging plasmid and 0.5 µg of pMD2.G envelope plasmid was transfected into the 293T cells using calcium phosphate transfection. In all, 48 hours after transfection, media containing virus was collected and filtered sterilized through 0.45 µm filters and flash frozen in 1 ml aliquots.

For lentiviral infection in 293T cell lines, $2.5 \times 10^5$ cells were seeded in six-well plates. 24 hours after initial seeding, 1 ml of virus (or media for mock infection) along with 2 ml of media supplemented with 10 µg/ml of polybrene was added to each well. The cells were washed with PBS and fed fresh media 24 h post-infection. Blasticidin or puromycin selection begun 48 h after infection at a concentration of 2 µg/ml. Fresh media supplemented with blasticidin or puromycin was added every other day and continued until the mock infection had no observable living cells. Stable integration and expression of each construct was verified via immunoblotting.

**Transient transfection and protein purification**

293T cells were transfected via calcium phosphate transfection method. Briefly, $2.5 \times 10^6$ cells were seeded on 10 cm$^2$ tissue culture grade plates followed by transfection with 10 µg of plasmid DNA (empty pcDNA3.1, pcDNA3.1-TRMT1-Strep or pcDNA3.1-Strep-TRMT1L). Cells were harvested 48 hours later by trypsin and neutralization with media, followed by centrifugation of the cells at 700 × g for 5 min followed by subsequent PBS wash and a second centrifugation step.

Protein was extracted by the Hypotonic Lysis protocol immediately after cells were harvested post-transfection. Cell pellets were resuspended in 0.5 mL of a hypotonic lysis buffer (20 mM HEPES pH 7.9, 2 mM MgCl2, 0.2 mM EGTA, 10% glycerol, 0.1 mM PMSF, 1 mM DTT) per plate. Cells were kept on ice for 5 min and then underwent three freeze-thaw cycles. NaCl was added to the extracts to a concentration of 0.4 M. After incubation on ice for 5 mins, the lysates were spun down at 14,000 × g for 15 min at 4 °C. The supernatant extract was removed, and an equal volume of Hypotonic Lysis buffer supplemented with 0.2% NP-40 was added to achieve the final whole cell lysates.

Purification of Strep-tagged proteins was carried out as previously described [13]. Briefly, cell lysates from the transiently-transfected cell lines were incubated with 50 µl of MagStrep "type3" XT beads (IBA Life Sciences) for two hours at 4 °C. Magnetic resin was washed three times in 20 mM

HEPES pH 7.9, 2 mM MgCl2, 0.2 mM EGTA, 10% glycerol, 0.1% NP-40, 0.2 M NaCl, 0.1 mM PMSF, and 1 mM DTT. Proteins were eluted with 1X Buffer BX (IBA LifeSciences) containing 10 mM D-biotin. Purified proteins fractionated on a 4-12% NuPAGE Bis-Tris polyacrylamide gel (ThermoFisher), stained with SYPRO Ruby Protein Gel Stain, and imaged on an Azure Sapphire Biomolecular Imager.

**In vitro methyltransferase assays**

DNA templates for *in vitro* transcription were generated by PCR. A list of double-stranded gBlock DNA (IDT) used as PCR templates are listed in Supplementary Table 3. Each template was designed with a T7 promoter upstream of the tRNA gene sequence. Each template was PCR amplified using a T7 Forward Primer (listed in Supplementary Table 3) and a specific reverse primer complementary to the 3' end of each tRNA species. PCR amplification was done using Herculase II Fusion DNA Polymerase (Agilent Technologies). The PCR parameters are as follows: 95 °C for 2 min followed by 35 cycles of 95 °C for 20 s, 48 °C for 20 s and 72 °C for 30 s, ending with 72 °C for 2 min. The PCR products were then resolved on a 2% agarose gel where bands were excised, and DNA was purified using the Qiagen Gel Extraction Kit. *In vitro* transcription was done using Optizyme T7 RNA Polymerase (ThermoFisher) following standard procedures. Reactions were incubated at 37 °C for 3 h followed by DNase treatment (RQ1 DNase, Promega) at 37 °C for 30 min. RNA was then purified using RNA Clean and Concentrator Zymo-Spin IC Columns (Zymo Research). tRNA transcripts were visualized on a 15% polyacrylamide, 7 M urea gel stained with SYBR Gold nucleic acid stain.

Prior to incubation with purified protein from human cells, the tRNA was first refolded by initial denaturation in 5 mM Tris pH 7.5 and 0.16 mM EDTA and heated to 95 °C for two minutes before a two-minute incubation on ice. Refolding was conducted at 37 °C for 20 min in the presence of HEPES pH 7.5, MgCl2, and NaCl. For each methyltransferase reaction, 100 ng of refolded tRNA was incubated with 25 nM of purified protein elution along with 50 mM TRIS pH 7.5, 0.1 mM EDTA, 1 mM DTT, 0.5 mM S-adenosylmethionine (NEB) for 6 h at 30 °C. RNA was purified using RNA Clean and Concentrator Zymo-

Spin IC Columns where the RNA was resuspended in 6 µl of RNase-free water. RNA samples were analyzed by LC-MS or primer extension as described above.

**Reporter assays**

The pcDNA3.1-10xArg-Nano reporter plasmid was previously described [13]. The 10xArg codon was replaced using BamH1-Kpn1 digestion and T4 ligation with double-stranded annealed oligonucleotide fragments encoding 10X tyrosine or serine codons (Supplemental Table 3). Plasmids were expressed in 293T cells by transient transfection using Lipofectamine 3000 (Invitrogen) and protein was extracted by above procedures. Immunoblotting against the FLAG tag was used to quantify nano-luciferase translation normalized by the internal firefly luciferase expression.

**Statistical analyses and reproducibility**

All statistics and graphs were performed and generated using GraphPad Prism software. Where applicable, error bars represent the standard deviation. Statistical tests and the number of times each experiment was repeated are stated in each figure legend.

**a.** >NG_054900.1:5828-17661 Homo sapiens tRNA methyltransferase 1 (TRMT1), RefSeqGene on chromosome 19

ccccaaaacgaggggatatcagcttaccgaggccgcaggttttcctagtccctacctcatagatatgtaggac
atccccgggcccggaatgcggctctctgaccctctctgtgccctcccccgcccccgaaccagGCTTGGCGGG
CGGAGGCGCCAGCGGATGTCTC**ATGCAAGGATCGTCTCTGTGGCTAAGCCTCACTTTCCGCTCCGCCCGGGTG**
M Q G S S L W L S L T F R S A R V
**CTCTCTAGAGCCCGGTTTTTCGAGTGGCAGTCTCCAGGGCTGCCGAATACAGCAGCGATGGAGAACGGCACCG**
L S R A R F F E W Q S P G L P N T A A M E N G T
**GGCCCTACGGAGAAGAACGTCCACGTGAAGTCCAGGAGACGACAGTCACCGAGGGGGCTGCCAAAATCGCCTT**
G P Y G E E R P R E V Q E T T V T E G A A K I A F
**TCCCAGTGCCAACGAGGTCTTTTATAACCCGGTGCAGGAATTCAATCGGGACCTGAC**gtgagcaggggtcaga
P S A N E V F Y N P V Q E F N R D L T
cgttagcctggcatcggctagtgggtcagtgtggatggcggg

**b.** >NC_000001.11:c185157537-185118101 Homo sapiens chromosome 1, GRCh38.p14 Primary Assembly

accgtagaaaccgaaccctcttacccaattggacgccgtccccagtacgcttctgcgcagtttgtggtggcgt
cGTAACGCCTCGGGGAAAGGGAAGCGGACGGGCATCTGGAATCGCTGCCTCTGGCTTTCTGTTTTCTACTAAC
AGGATTTGGTCACTGGTTCTTCATCTTTTGTCTGTTGCACGCATCCCGCCCTCCCCACTTGCTTCCCCACTCC
TTGGATCCAGCCCTGTGGGCATTCACGTCAGTTCTCTGACCCCGCCGTGAGCCCCGCTCCGGGTCCCCGGGCG
GGCTTGGCACGGAGGCGGTAACT**ATGGAGAATATGGCGGAGGAGGAGCTGCTGCCCCTGGAGAAGGAGGAGGT**
M E N M A E E E L L P L E K E E V
**GGAGGTGGCCCAGGTCCAGGTCCCGACCCCGGCCCGGGACTCGGCTGGGGTCCCAGCTCCGGCCCCGGATTCG**
E V A Q V Q V P T P A R D S A G V P A P A P D S
**GCTCTGGACTCGGCTCCGACTCCGGCCTCGGCTCCAGCCCCAGCCCCTGCCCTGGCCCAGGCTCCGGCCCTGT**
A L D S A P T P A S A P A P A P A L A Q A P A L
**CCCCGTCCCTAGCCTCTGCCCCTGAGGAGGCTAAAAGCA**gtaagtgcagaaggcccagatctttctgctgcag
S P S L A S A P E E A K S
aagagagaaagtgcgccttgctgggaagtaggggaggcccttcaccgggatggttttatggggcaaggcaggt
ttaggaaaatggtggggaggaa

**Supplemental Figure S1.** Genomic sequences of (a) TRMT1 and (b) TRMT1L. Intronic sequences in lowercase, exon sequences in uppercase, open reading frame in bold, underlined sequences are targeted by CRISPR guide RNAs, and sequences deleted in TRMT1-KO or TRMT1L-KO cell lines are highlighted in yellow.

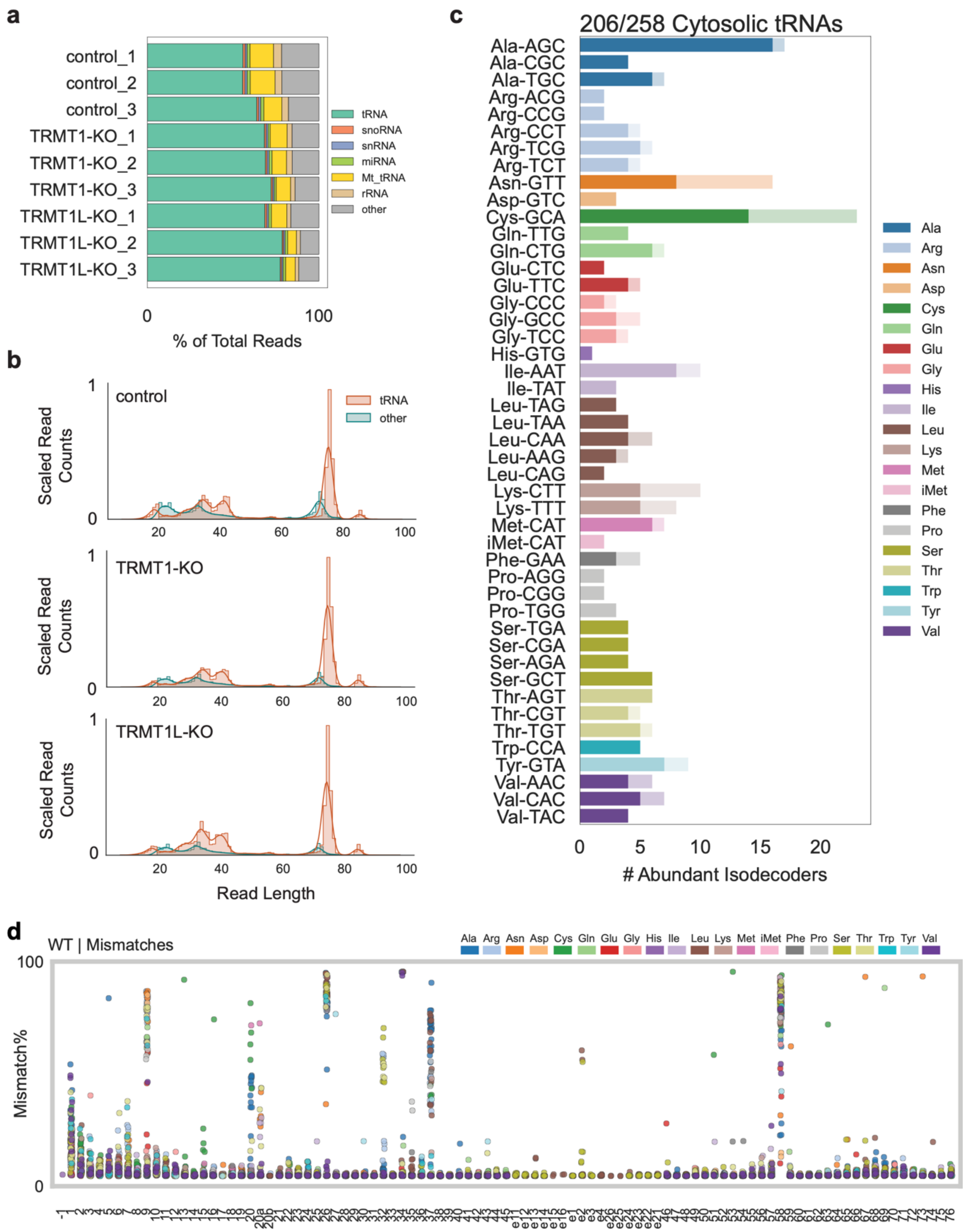


**Supplemental Figure S2.** Sequence coverage of tRNAs using the OTTR-Seq approach. (a) Percentage of total reads mapped to the indicated RNA classes for each sample. (b) Scaled read counts for tRNA versus other RNAs at each read length. (c) Coverage of cytosolic tRNA isodecoders. Shaded portion of each bar represents the tRNA isodecoders that were detected within the tRNA isodecoder family (d) Mismatch incorporation percentage at each position for tRNA isoacceptors in control-WT samples.

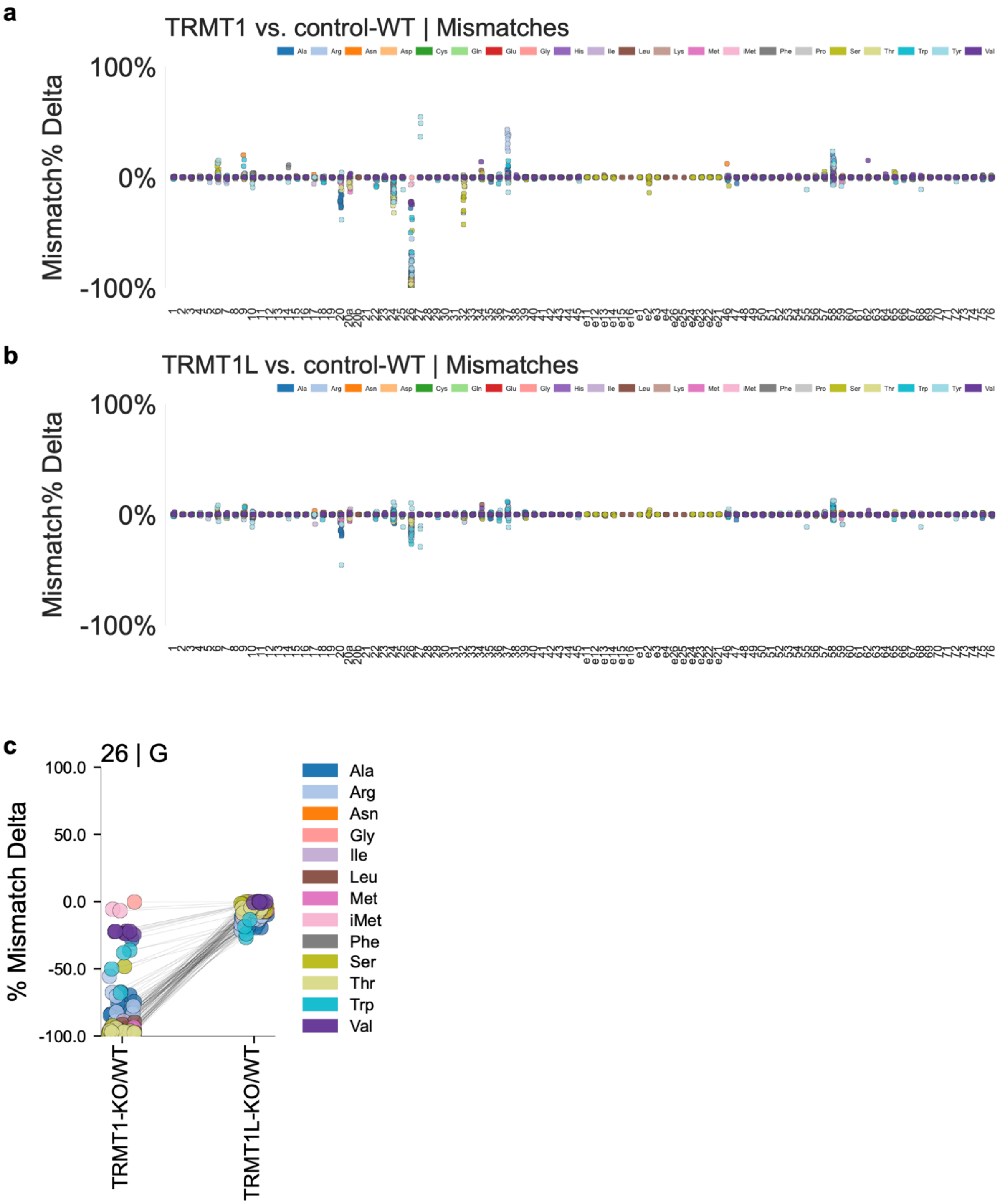


**Supplemental Figure S3.** Misincorporation frequency changes in TRMT1-KO and TRMT1L-KO cell lines compared to the control-WT cell line. (a) Change in mismatch incorporation percentage at each position for tRNA isoacceptors in TRMT1-KO versus control-WT cell lines. (b) Change in mismatch incorporation percentage at each position for tRNA isoacceptors in TRMT1L-KO versus control-WT cell lines. (c) Change in mismatch incorporation percentage at position 26 for each tRNA isoacceptor in the TRMT1-KO or TRMT1L-KO cell line relative to the control-WT cell line.

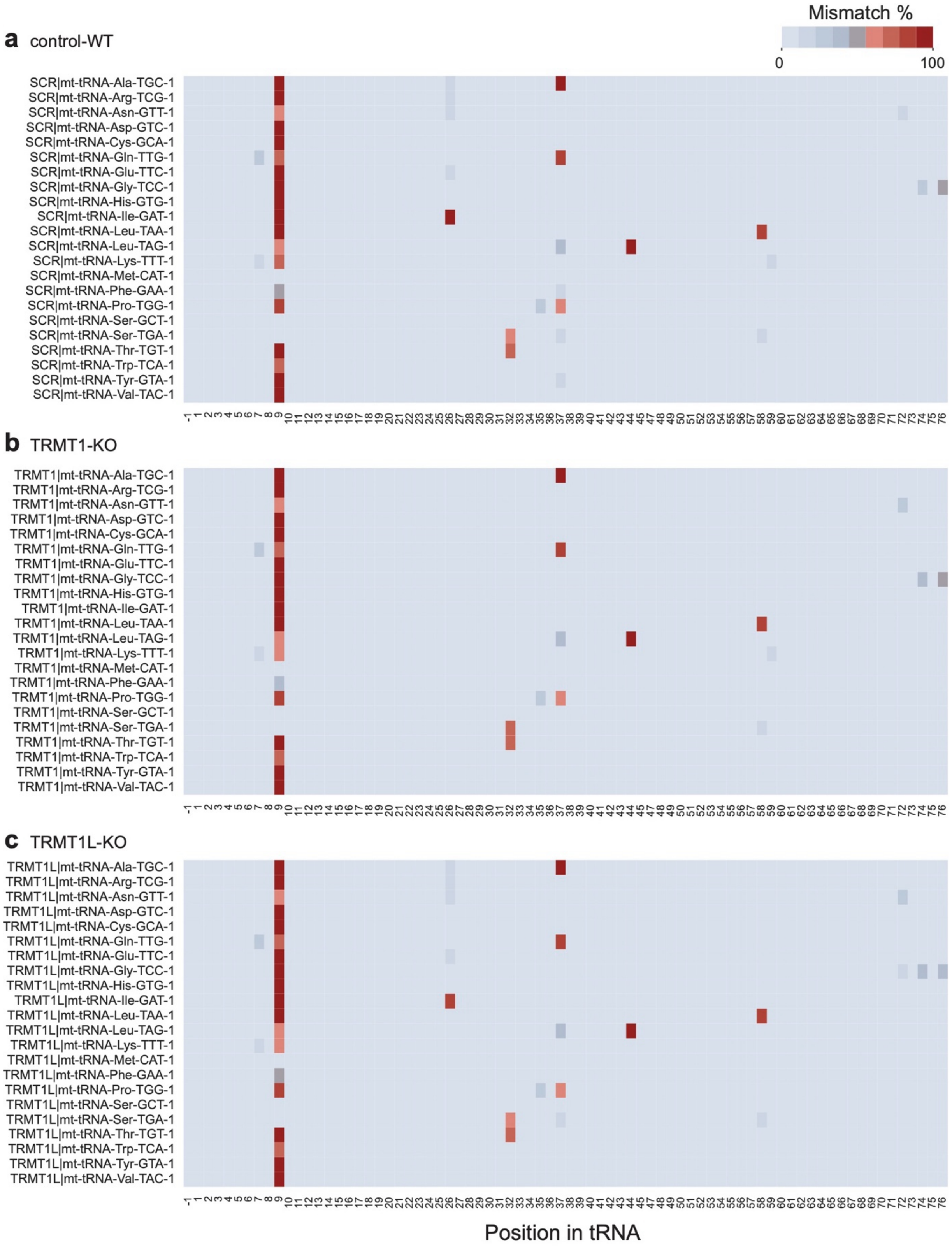


**Supplemental Figure S4.** Mismatch percentage heatmap of mitochondria (mt)-tRNAs from (a) control-WT, (b) TRMT1-KO, and (c) TRMT1L-KO cell lines.

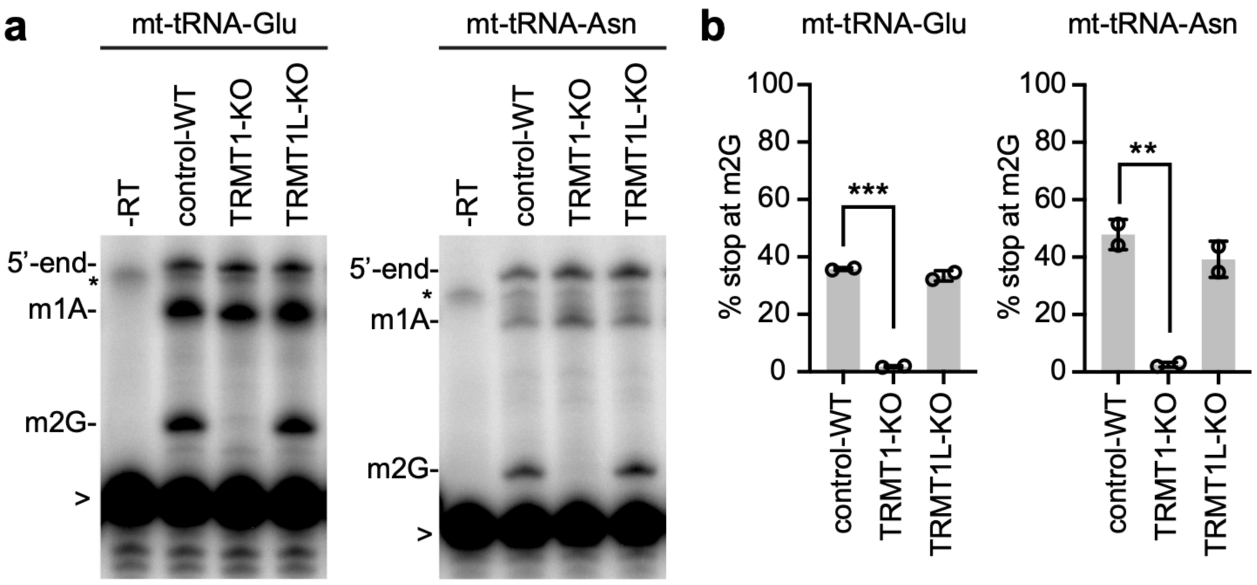


**Supplemental Figure S5.** TRMT1 is required for m2G formation in mt-tRNA-Glu and mt-tRNA-Asn. (a) Primer extension analysis of mt-tRNA-Glu and mt-tRNA-Asn harvested from the indicated human cell lines. (b) Quantification of m2G formation in mt-tRNA-Glu and mt-tRNA-Asn. Bars represent the standard deviation from the mean. Statistical analysis was performed using one-way ANOVA and significance calculated using Dunnett's multiple comparison test. ***$P \leq 0.001$, **$P \leq 0.01$. *$P \leq 0.05$.

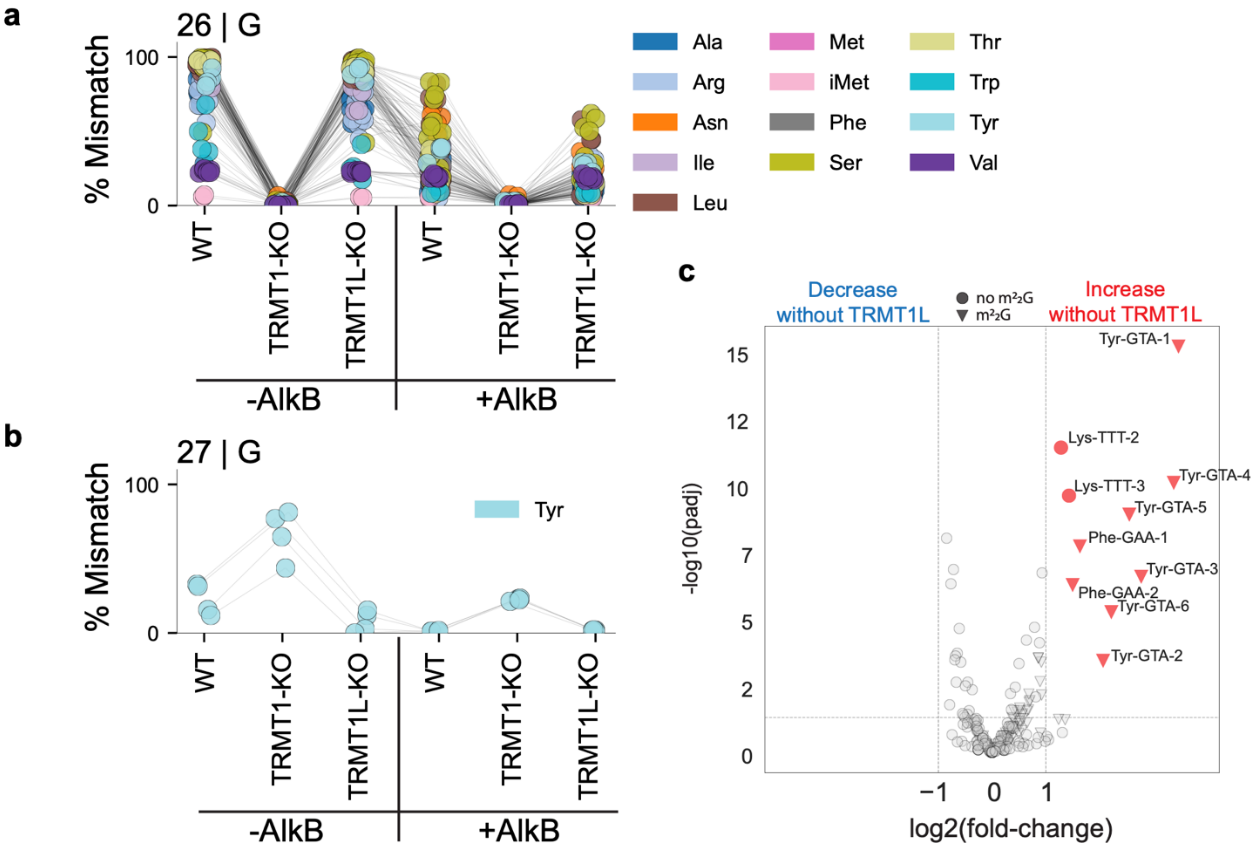


**Supplemental Figure S6.** Mismatch incorporation frequency in control-WT, TRMT1-KO, and TRMT1L-KO cell lines. (a) Mismatch incorporation percentage at position 26 for each tRNA isoacceptor in the control-WT, TRMT1-KO, and TRMT1L-KO cell lines. (b) Mismatch incorporation percentage at position 27 for tRNA-Tyr-GUA isodecoders in the control-WT, TRMT1-KO, and TRMT1L-KO cell lines. For (a) and (b), OTTR-Seq was performed with untreated RNA (-AlkB) or with RNA incubated with AlkB (+AlkB). (c) Volcano plot of tRNA levels in TRMT1L-KO versus control-WT cell lines. Triangles represent tRNAs that are modified with m2,2G at position 26. Circles represent tRNAs without m2,2G. Red and blue denote tRNAs that are increased and decreased, respectively, in the TRMT1L-KO cell lines.

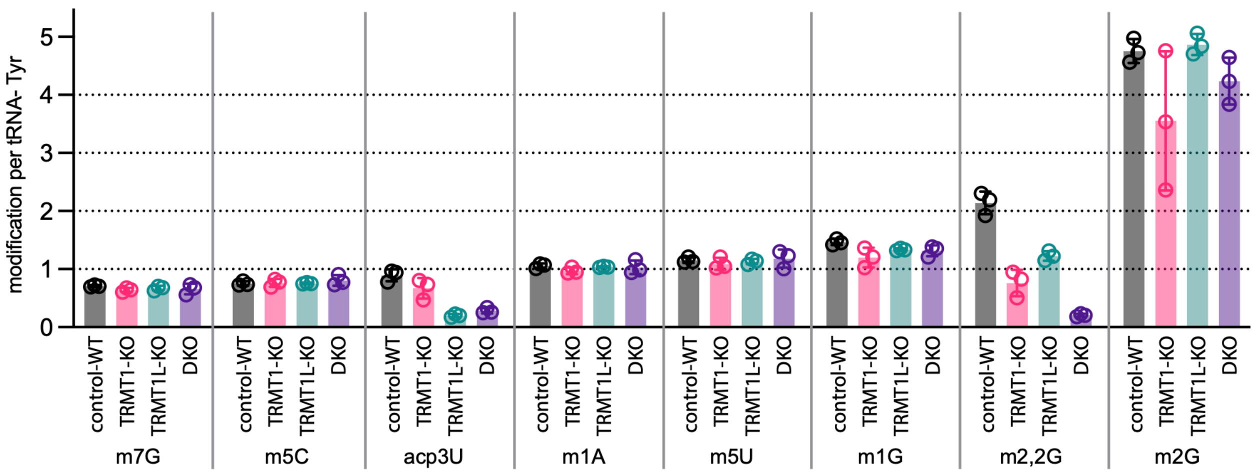


**Supplemental Figure S7.** LC-MS analysis of nucleosides with stable isotope-labeled internal standards (SILIS). Graph represents the modifications per purified tRNA-Tyr from the indicated human 293T cell lines.

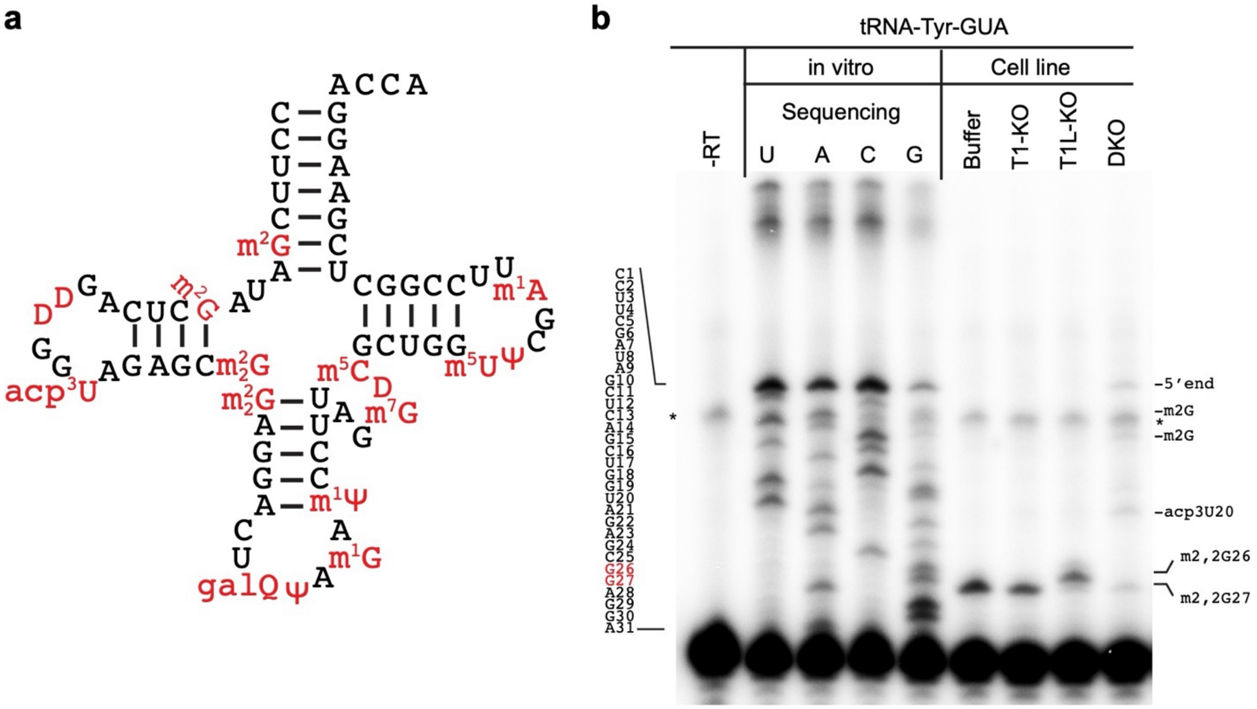


**Supplemental Figure S8.** Primer extension analysis of human tRNA-Tyr-GUA. (a) Secondary structure of human tRNA-Tyr with modifications noted. (b) Primer extension gel of in vitro transcribed tRNA-Tyr or total RNA samples isolated from the indicated cell lines. Sequencing ladder is noted on the left. -RT is a control reaction without reverse transcriptase.

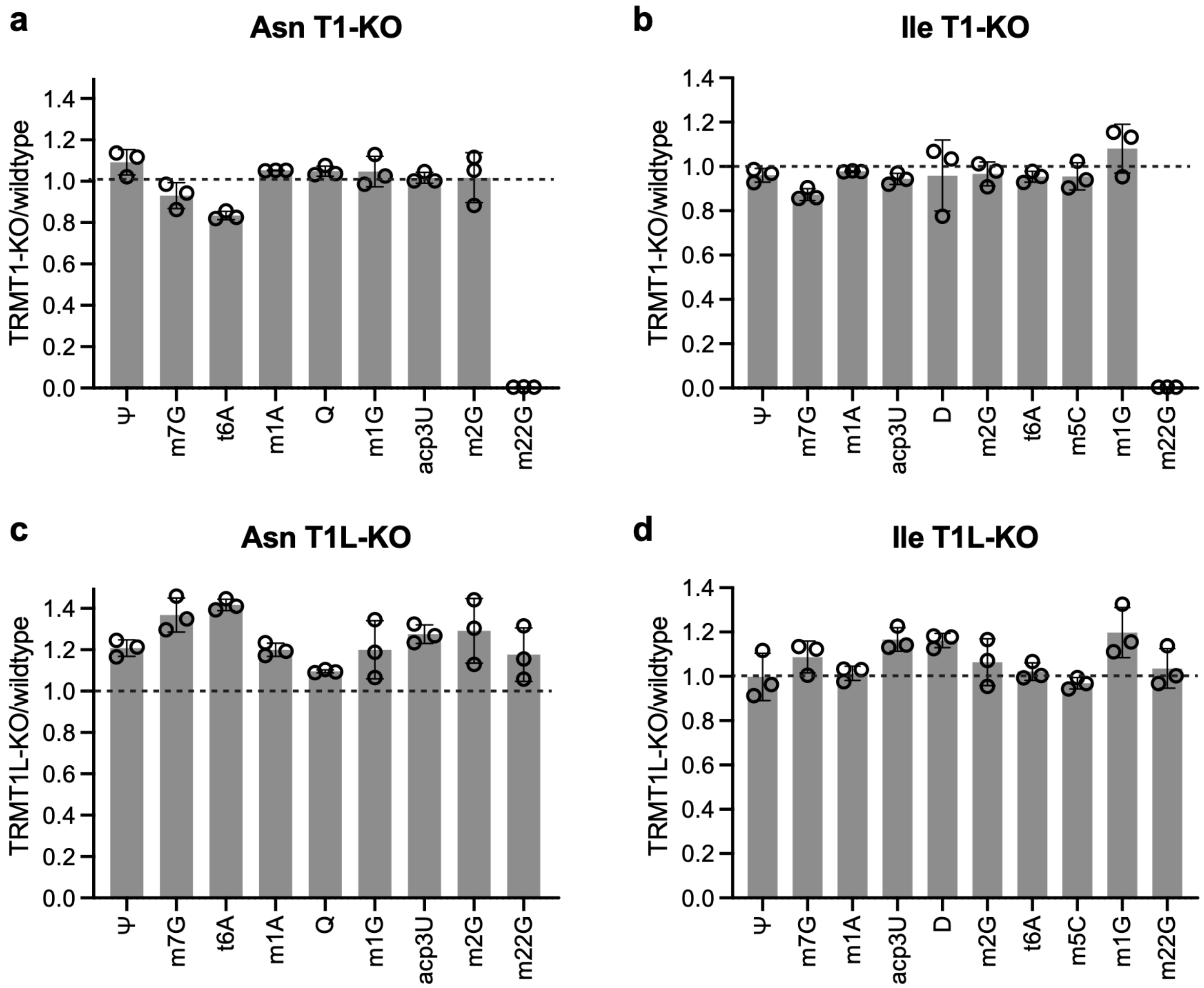


**Supplemental Figure S9.** LC-MS analysis of tRNAs isolated from TRMT1-KO and TRMT1L-KO cell lines. (a, b) Relative peak intensity values of the indicated modifications detected by LC-MS in purified: (a) tRNA-Asn or (b) tRNA-Ile from the TRMT1-KO cell line versus control-WT cell line. (c, d) Relative peak intensity values of the indicated modifications detected by LC-MS in purified: (c) tRNA-Asn or (d) tRNA-Ile from the TRMT1L-KO cell line versus control-WT cell line. (f) Relative peak intensity values of the indicated modifications detected by LC-MS in purified tRNA-Ser from the TRMT1-KO cell line versus control-WT cell line.

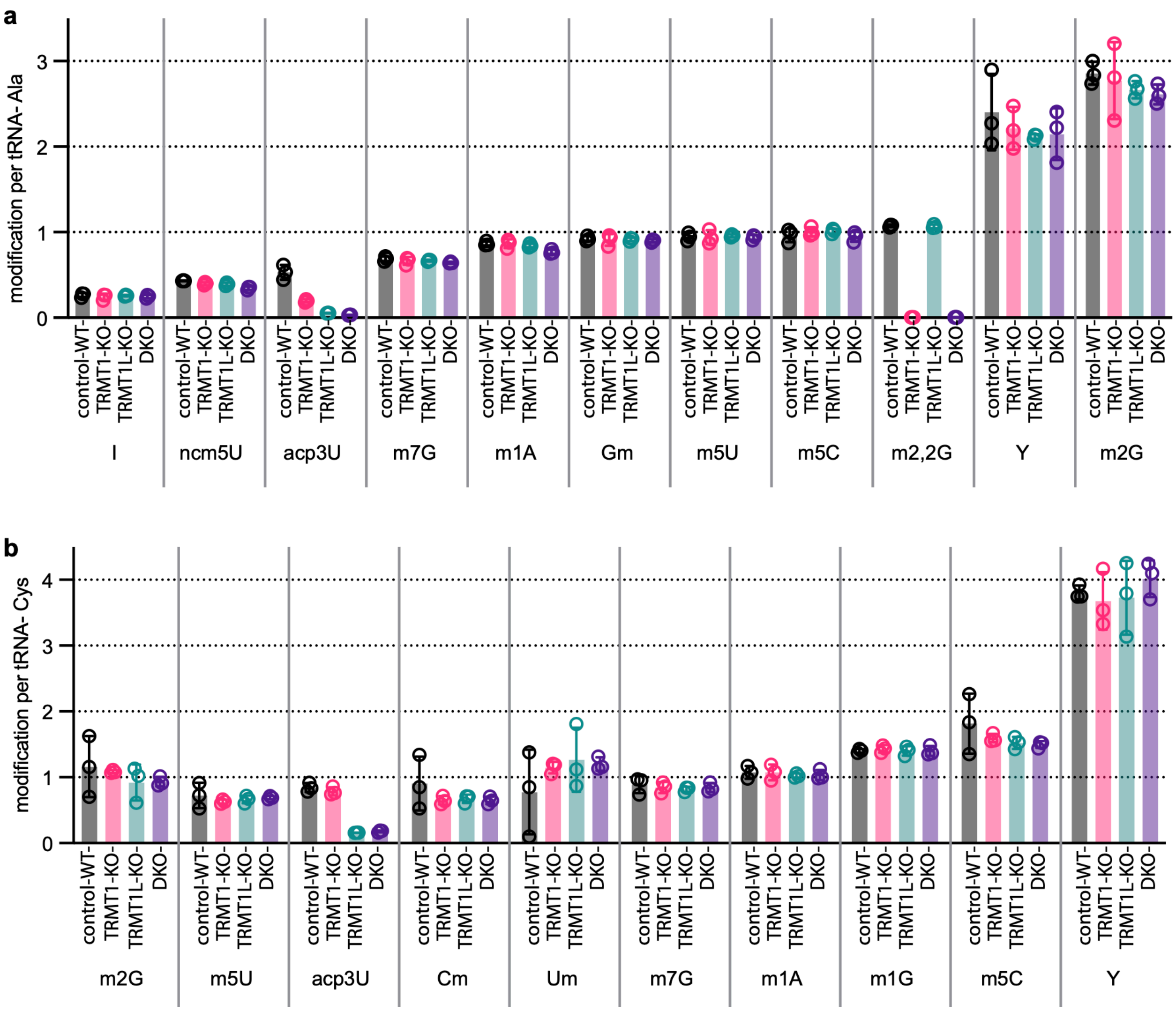


**Supplemental Figure S10.** LC-MS analysis of nucleosides with stable isotope-labeled internal standards (SILIS). (a) Modifications per purified tRNA-Ala from the indicated human 293T cell lines. (b) Modifications per purified tRNA-Cys from the indicated human 293T cell lines. I, inosine. Y, pseudouridine.

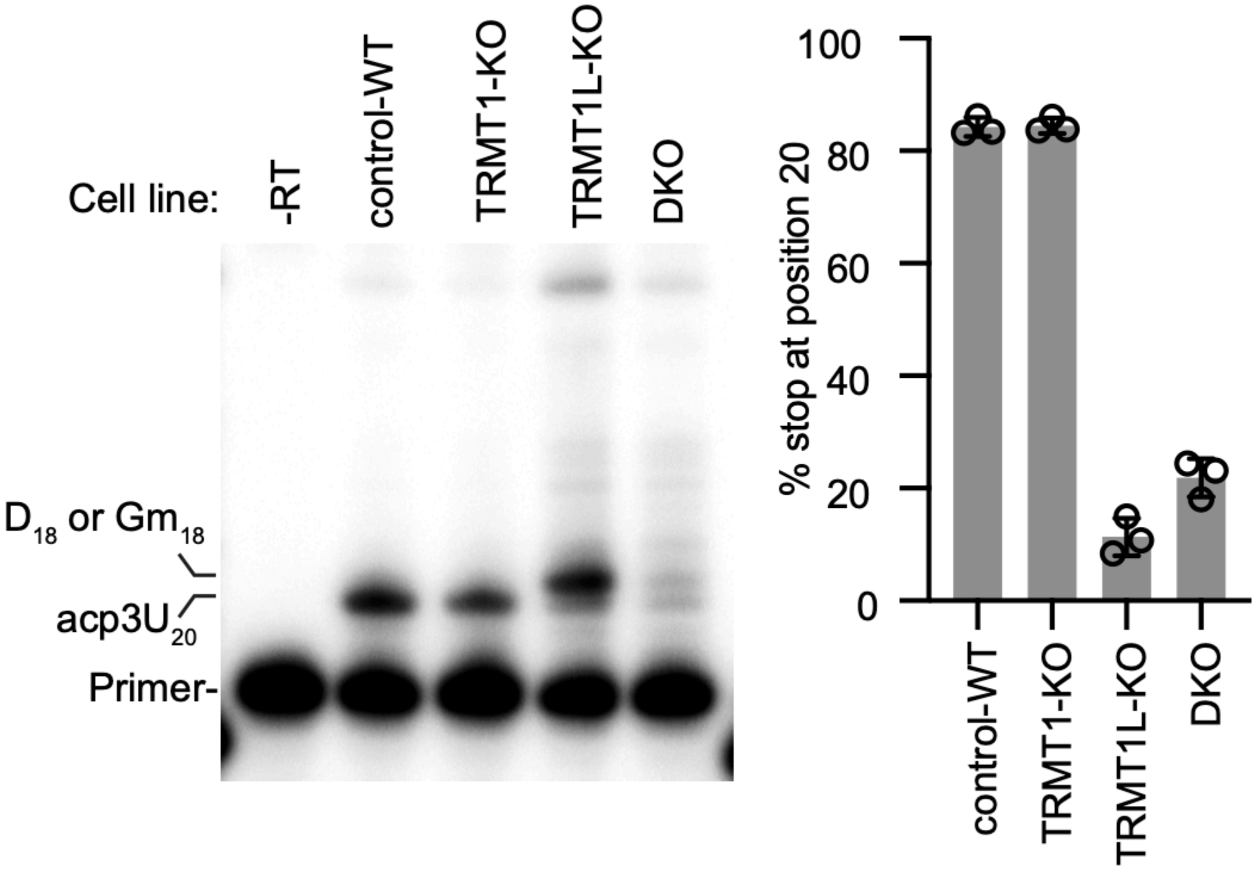


**Supplemental Figure S11.** Representative primer extension gel of tRNA-Cys from the indicated 293T cell lines. -RT represents a reaction performed without reverse transcriptase. Quantification of the stop signal relative to readthrough plus stop signal is noted to the right.

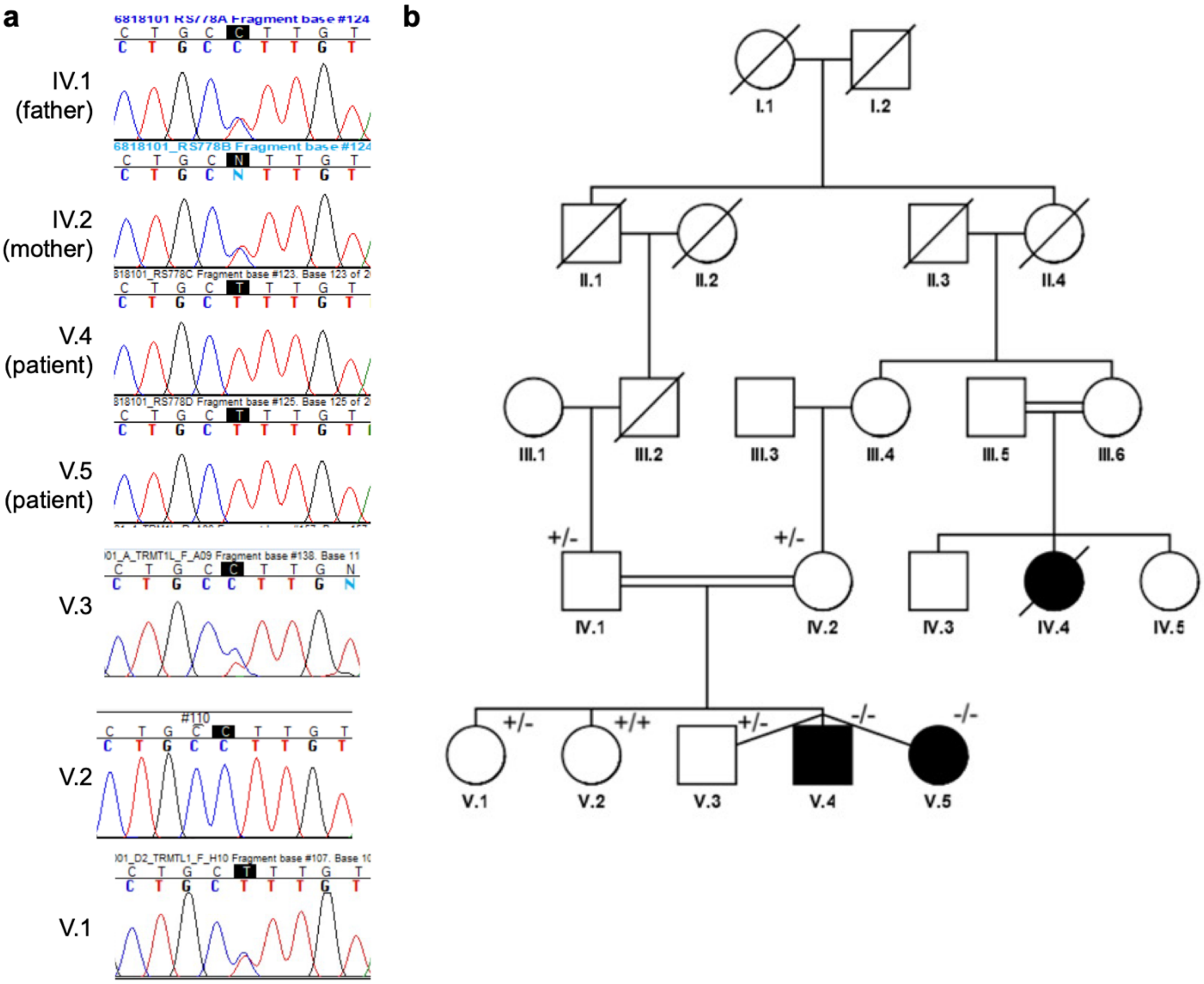


c

| Analysis | SIFT | PolyPhen | Mutation Assessor | Mutation taster |
|---|---|---|---|---|
| Score | 0.01 | 0.94 | 0.797 | 80\|20 |
| Predicted outcome | Deleterious | Probably damaging | Medium impact | Deleterious |

**Supplemental Figure S12.** Identification of a pathogenic variant in the human *TRMT1L* gene. (a) Sequencing chromatograms from the indicated individuals from the pedigree in (b). The patients with the homozygous variant are patients V.4 and V.5. (c) Scores and predicted outcomes for the P512L variant based upon the indicated pathogenicity prediction algorithms.

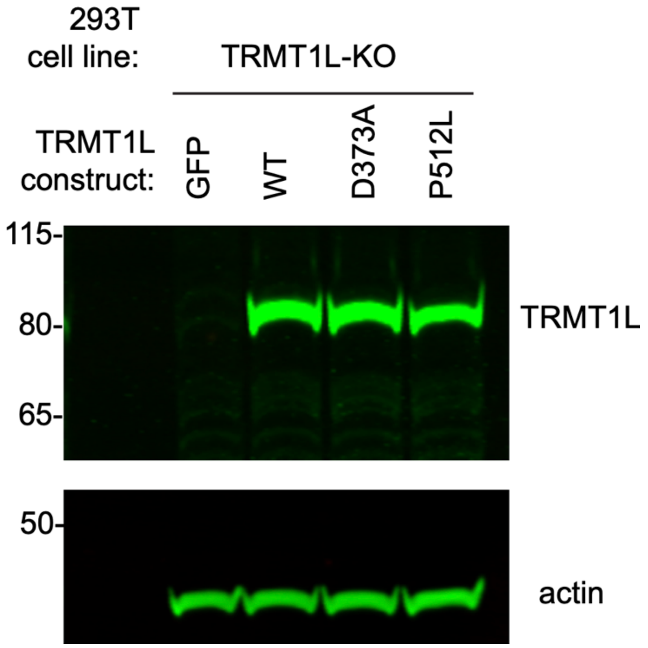


**Supplemental Figure S13.** Immunoblot analysis of the indicated TRMT1L-KO cell lines containing a lentiviral-integrated construct. The blot was probed with anti-TRMT1L or anti-actin antibodies.

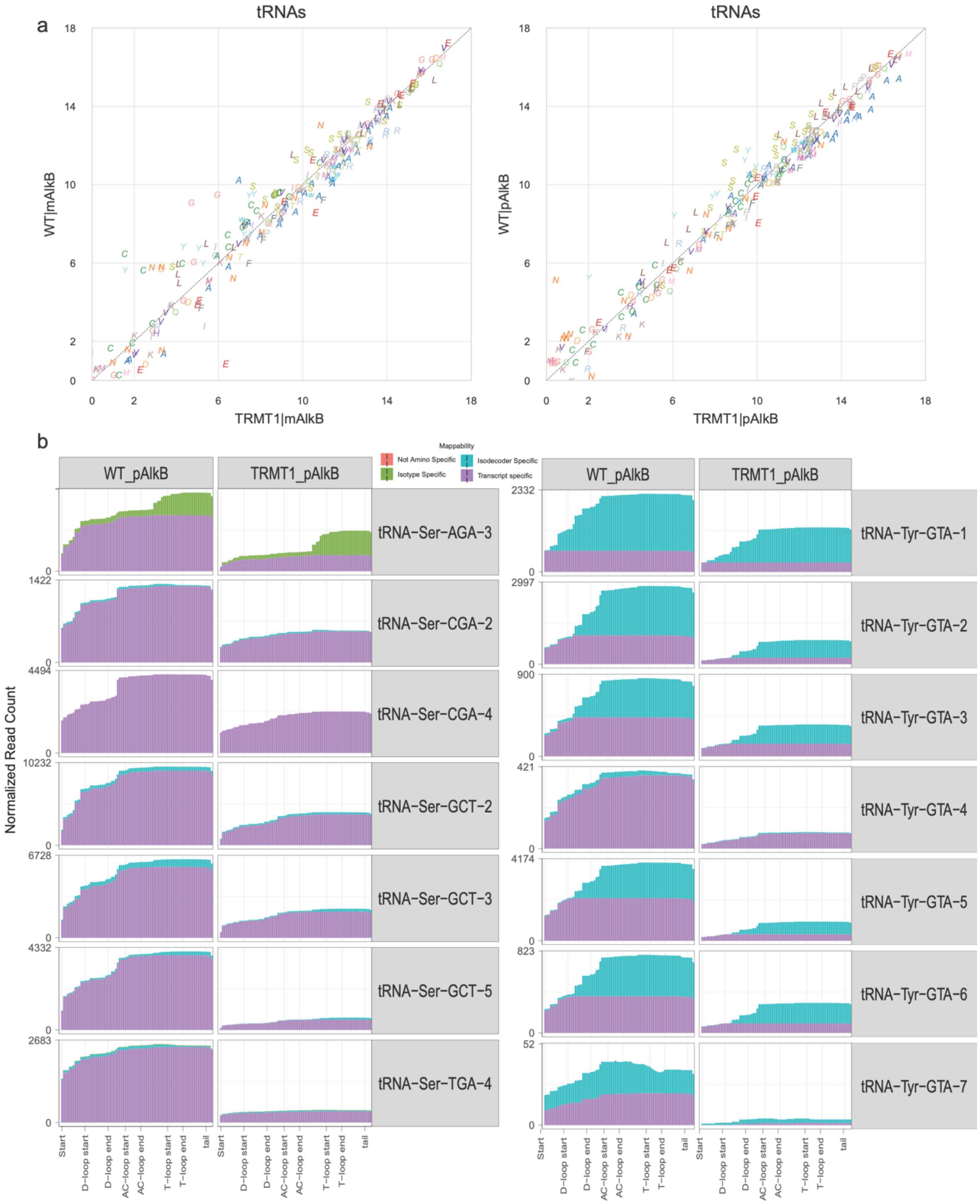


**Supplemental Figure S14.** Changes in full-length tRNA levels in the TRMT1-KO cell line. (a) Total read counts of the indicated full-length tRNAs in the TRMT1-KO cell line versus the control-WT cell line without or with AlkB. (b). Raw normalized read counts or tRNA-Ser and tRNA-Tyr isodecoders that decreased in the TRMT1-KO cell line.

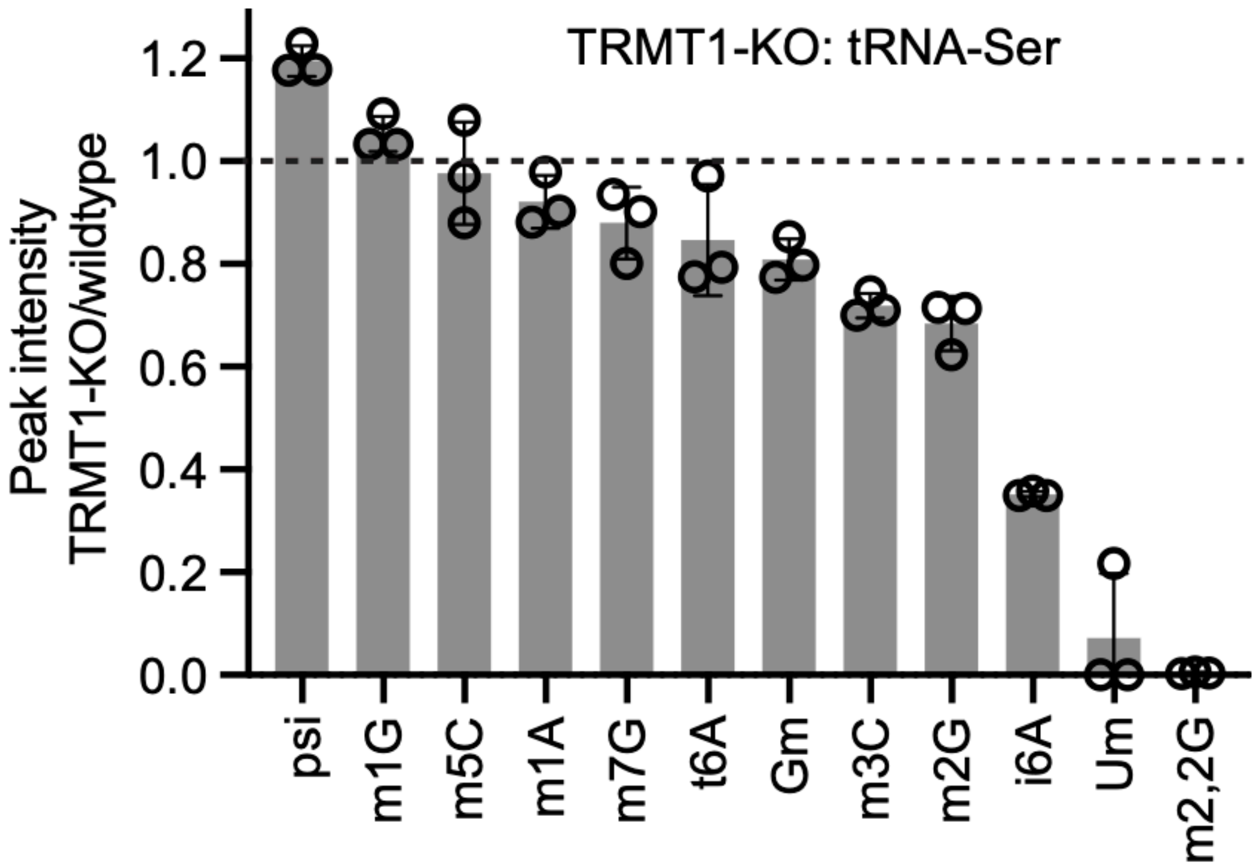


**Supplemental Figure S15.** Relative peak intensity of nucleosides detected by LC-MS in purified in tRNA-Ser from TRMT1-KO cells relative to control-WT cells.

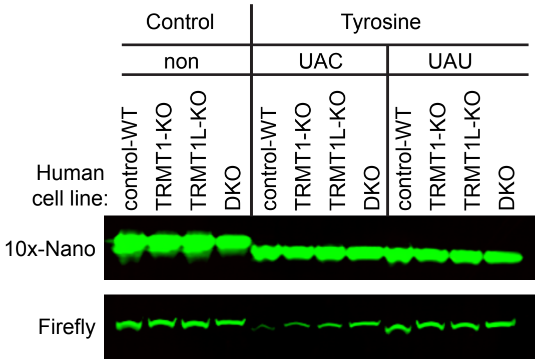


**Supplemental Figure S16.** Representative immunoblot of nanoluciferase codon reporter assay. Source data contains additional immunoblots for each replicate.

| Isotype | Anticodon | Isodecoders | Modification | Fraction of Isodecoders |
|---|---|---|---|---|
| Ala | IGC | 1-15,24 | $m^2{}_2G$ | 16/16 |
| Ala | CGC | 1,2,3,4 | $m^2{}_2G$ | 4/4 |
| Ala | UGC | 1,2,3,4,5 | $m^2{}_2G$ | 5/6 |
| Arg | ICG | 1,2 | $m^2{}_2G$ | 2/2 |
| Arg | CCG | 1 | $m^2{}_2G$ | 1/2 |
| Arg | UCG | 1,2,3,4,5 | $m^2{}_2G$ | 5/5 |
| Arg | UCU | 4 | $m^2{}_2G$ | 1/4 |
| Asn | GUU | 1,2,4,5,6,9 | $m^2{}_2G$ | 6/7 |
| Ile | AAU | 2,3,4,5,6,7,8,12 | $m^2{}_2G$ | 8/8 |
| Ile | UAU | 1,2,3 | $m^2{}_2G$ | 3/3 |
| iMet | CAU | 1,2 | $m^2G$ | 2/2 |
| Leu | AAG | 1,2,3 | $m^2{}_2G$ | 3/3 |
| Leu | CAA | 1,2,3,4 | $m^2{}_2G$ | 4/4 |
| Leu | CAG | 1,2 | $m^2{}_2G$ | 2/2 |
| Leu | UAA | 1,2,3,4 | $m^2{}_2G$ | 4/4 |
| Leu | UAG | 1,2,3 | $m^2{}_2G$ | 3/3 |
| Met | CAT | 1,2,3,4,5,6 | $m^2{}_2G$ | 6/6 |
| Phe | GAA | 1,2 | $m^2{}_2G$ | 2/2 |
| Ser | AGA | 1,2,3,4 | $m^2{}_2G$ | 4/4 |
| Ser | CGA | 1,2,3,4 | $m^2{}_2G$ | 4/4 |
| Ser | GCU | 1,2,3,4,5 | $m^2{}_2G$ | 5/5 |
| Ser | UGA | 1,2,3,4 | $m^2{}_2G$ | 4/4 |
| Thr | AGU | 1,2,3,4,5,6 | $m^2{}_2G$ | 6/6 |
| Thr | CGC | 1,2,3,4 | $m^2{}_2G$ | 4/4 |
| Thr | UGU | 1 | $m^2{}_2G$ | 1/4 |
| Trp | CCA | 1,2,3,4 | $m^2{}_2G$ | 4/4 |
| Tyr | GUA | 1,2,3,4,5,6 | $m^2{}_2G$ | 6/6 |
| Val | AAC | 1,2,3,4,5 | $m^2G$ | 5/5 |
| Val | CAC | 1,2,4,5 | $m^2G$ | 4/5 |
| Val | CAC | 1,2 | $m^2G$ | 2/4 |

Table 1: Cytosolic tRNAs modified at position 26 by TRMT1.

| Isotype | Anticodon | Modification |
|---|---|---|
| mt-Ala | UGC | $m^2G$ |
| mt-Arg | UCG | $m^2G$ |
| mt-Asn | GUU | $m^2G$ |
| mt-Glu | UUC | $m^2G$ |
| mt-Ile | GAU | $m^2{}_2G$ |

Table 2: Mitochondrial tRNAs modified at position 26 by TRMT1.

**Supplemental Table S3.** DNA Oligonucleotides and antibodies

| **Primer extension** | Sequence (5'-> 3') |
|---|---|
| mt-tRNA-Glu | GGACTACAACCACGACCAATGATATGAA |
| mt-tRNA-Asn | TAAACCCACAAACACTTAGTTAACAGC |
| Tyr-GUA_58-33 | TCGAACCAGCGACCTAAGGATCTACA |
| pre-tRNA-Tyr-R-PE | GAATCGAACCAGCGACCTAAGGAT |
| Ile-AAT-PE | GAACCCGCGACCTTGGCGTTATTA |
| mt-Ile-GAU | TTAAGCTCCTATTATTTACTCTATC |
| Met-CAU | AACTCACGACCTTCAGATTATG |
| Cys-GCA | GGGACCTCTTGATCTGCAGTCAAATG |
| **tRNA purification** | |
| tRNA-Ala | /5BiosG/TGCCGGGGATYGAACCCGGGRCCTC |
| tRNA-Ser-AGA | /5BiosG/CGTAGTCGGCAGGATTCGAACCTGCG |
| tRNA-Ser-AGA | /5BiosG/CGTAGTCGGCAGGATTCGAACCTGCG |
| tRNA-Ser-CGA | /5BiosG/GARCAGGATTYGAACCTGCGCGGGGA |
| tRNA-Ser GCT | /5BiosG/CGACGAGGATGGGATTCGAACCCACG |
| tRNA-Arg | /5BiosG/AGGASTCGAACCYDSAATCTTCTGAT |
| tRNA-Tyr | /5BiosG/AAATGGTCCTTCGAGCCGGAWTCGAACCAGCGA |
| tRNA-Cys | /5BiosG/AGGGGGCACCYGGATTTGAACCRGGGAC |
| tRNA-Ile-TAT | /5BiosG/TGCTCCAGGTGAGGMTCGAACTCACAAC |
| tRNA-Ile-AAT | /5BiosG/TGGCCMGTACGGGGATCGAACCCGCGAC |
| tRNA-Asn | /5BiosG/GTCCCTGGGTGGGMTCGAACCACCAAC |
| **Northern-blot** | |
| tRNA-Tyr-PHA(Dloop) | CTA CAG TCC TCC GCT CTA CC |
| tRNA-Tyr-Tloop | GACCTAAGGATCTACAGTCCT |
| tRNA-Ser | GACGARGRTGGGATTCGAACCCA |
| Ala-AGC-8-PHA (Dloop) | AGCGAGCGCTCTACCATTTG |
| Ala-AGC-8-Tloop | GGAGGATGCGGGCATCGATC |
| tRNA-Lys-UUU | GGG ACT TGA ACC CTG GAC CCT |
| U6 snRNA | CGTTCCAATTTTAGTATATGTGCTGCCGAAGCGA |
| Glu-TTC | TTCCCTGGCCGGGAATCG |
| 5' Tyr-GTA | CCAACTGAGCTATCGAAGG |
| 3' Tyr-GTA | TGGTCCTTCGAGCCGG |
| ac region Tyr-GTA | CGAACCAGCGACCTAAGGAT |
| **Cloning- reporter assay** | |
| 10X random-F | GATCC GGCCAGGGCCCTTGCGGCCAGGGCGGCTGC GGTAC |
| 10X random-F | C GCAGCCGCCCTGGCCGCAAGGGCCCTGGCC G |
| Tyr-TAC-F | GATCC TACTACTACTACTACTACTACTACTACTAC GGTAC |
| Tyr-TAC-R | C GTAGTAGTAGTAGTAGTAGTAGTAGTAGTA G |
| Tyr-TAT-F | GATCC TATTATTATTATTATTATTATTATTATTAT GGTAC |
| Tyr-TAT-R | C ATAATAATAATAATAATAATAATAATAATA G |
| Ser-TCT-F | GATCCTCTTCTTCTTCTTCTTCTTCTTCTTCTTCTGGTAC |
| Ser-TCT-R | CAGAAGAAGAAGAAGAAGAAGAAGAAGAAGAG |
| Ser-TCA-F | GATCCTCATCATCATCATCATCATCATCATCATCAGGTAC |
| Ser-TCA-R | CTGATGATGATGATGATGATGATGATGATGAG |
| Ser-TCC-F | GATCCTCCTCCTCCTCCTCCTCCTCCTCCTCCTCCGGTAC |
| Ser-TCC-R | CGGAGGAGGAGGAGGAGGAGGAGGAGGAGGAG |
| Ser-TCG-F | GATCCTCGTCGTCGTCGTCGTCGTCGTCGTCGTCGGGTAC |
| Ser-TCG-R | CCGACGACGACGACGACGACGACGACGACGAG |

| | |
|---|---|
| Ser-AGT-F | GATCCAGTAGTAGTAGTAGTAGTAGTAGTAGTAGTGGTAC |
| Ser-AGT-R | CACTACTACTACTACTACTACTACTACTG |
| Ser-AGC-F | GATCCAGCAGCAGCAGCAGCAGCAGCAGCAGCAGCGGTAC |
| Ser-AGC-R | CGCTGCTGCTGCTGCTGCTGCTGCTGCTGCTG |
| **Cloning- Mutagenesis** | |
| TRMT1 D233A-F | ATCGATCTGGCCCCCTATGGCAGC |
| TRMT1 D233A-B | GCTGCCATAGGGGGCCAGATCGAT |
| TRMT1L D373A-F | TTCATACATCTAGCCCCTTTTGGAACATCAGTG |
| TRMT1L D373A-R | CACTGATGTTCCAAAAGGGGCTAGATGTATGAA |
| **Cloning- CRISPR-edit** | |
| TRMT1 gPCR F6 | TTTCTGTGGCTACGAGAGCA |
| TRMT1 gPCR R6 | CCCAGGCAGGGAGATAAACTT |
| TRMT1L gs F2 | CACCGTCCGGCCCCGGATTCGGCTC |
| TRMT1L gs R2 | AAACGAGCCGAATCCGGGGCCGGAC |
| TRMT1L gs F5 | CACCGGGTAACTATGGAGAATATGG |
| TRMT1L gs R5 | AAACCCATATTCTCCATAGTTACCC |
| **Cloning- CRISPR-edit** | |
| TRMT1 gPCR F6 | TTTCTGTGGCTACGAGAGCA |
| TRMT1 gPCR R6 | CCCAGGCAGGGAGATAAACTT |
| TRMT1L gs F2 | CACCGTCCGGCCCCGGATTCGGCTC |
| TRMT1L gs R2 | AAACGAGCCGAATCCGGGGCCGGAC |
| TRMT1L gs F5 | CACCGGGTAACTATGGAGAATATGG |
| TRMT1L gs R5 | AAACCCATATTCTCCATAGTTACCC |
| **In vitro transcription** | |
| tRNA-Tyr-GTA gBlock | CAGAAGGAATTCGCTGCAGTAATACGACTCACTATAGCTTCGATAGCTCAGC TGGTAGAGCGGAGGACTGTAGATCCTTAGGTCGCTGGTTCGATTCCGGCTC GAAGCACCAGGATCCCGATGCA |
| tRNA-Tyr-GTA-gBlock G26A | CAGAAGGAATTCGCTGCAGTAATACGACTCACTATAGCTTCGATAGCTCAGC TGGTAGAGCAGAGGACTGTAGATCCTTAGGTCGCTGGTTCGATTCCGGCTC GAAGCACCAGGATCCCGATGCA |
| tRNA-Tyr-GTA-gBlock G27A | CAGAAGGAATTCGCTGCAGTAATACGACTCACTATAGCTTCGATAGCTCAGC TGGTAGAGCGAAGGACTGTAGATCCTTAGGTCGCTGGTTCGATTCCGGCTC GAAGCACCAGGATCCCGATGCA |
| tRNA-Tyr-GTA-1-1-Fx | GCAGTAATACGACTCACTATAGCTTCGATAGCTCAGCTG |
| tRNA-Tyr-GTA-1-1-Rx | TGGTGCTTCGAGCCGGAATCGAACC |
| tRNA-Ile-AAT-2-1-U27G gBlock | CAGAAGGAATTCGCTGCAGTAATACGACTCACTATAGGCCGGTTAGCTCAGT TGGTTAGAGCGGGGTGCTAATAACGCCTAGGTCGCGGGTTCGATCCCCGTA CTGGCCACCAGGATCCCGATGCA |
| Pre-tRNA-Tyr-5-1 gBlock | CAGAAGGAATTCGCTGCAGTAATACGACTCACTATAGCTTCGATAGCTCAGC TGGTAGAGCGGAGGACTGTAGCTACTTCCTCAGCAGGAGACATCCTTAGGT CGCTGGTTCGATTCCGGCTCGAAGCACCAGGATCCCGATGCA |
| Ile-AAT-PCR-R | TGGTGGCCAGTACGGGGATCGAAC |
| pre-tRNA-Tyr-R-PE | GAATCGAACCAGCGACCTAAGGAT |
| tRNA-Ile-AAT-2-1 gBlock | CAGAAGGAATTCGCTGCAGTAATACGACTCACTATAGGCCGGTTAGCTCAGT TGGTTAGAGCGTGGTGCTAATAACGCCAAGGTCGCGGGTTCGATCCCCGTA CTGGCCACCAGGATCCCGATGCA |
| T7 promoter | |
| **Antibody** | |

| | |
|---|---|
| TRMT1-G3 | Santa Cruz Biotechnology, sc-373687, 1: 500 dilution |
| TRMT1L | Abnova, H00081627-B01P, 1:1000 dilution |
| Flag-M2 | Sigma-Aldrich, F1804, 1:3333 dilution |
| Twin-Strep | Genscript, cat. No. A01732, 1:1000 dilution |
| Actin C4 | EMD Millipore, cat. No MAB1501, 1:2500 dilution |
| IRDye 800CW goat anti-mouse IgG | Thermofisher, 925-3221, 1:10000 dilution |

**Supplemental Table S4.**

| Compound Group | Compound Name | ISTD? | Precursor Ion | MS1 Res | Product Ion | Ret Time (min) | Delta Ret Time | Fragmentor | Collision Energy |
|---|---|---|---|---|---|---|---|---|---|
| A | A | Target | 268.1 | Wide | 136 | 5.239 | 1 | 180 | 21 |
| A SILIS | A SILIS | ISTD | 283 | Wide | 146 | 5.237 | 1 | 180 | 21 |
| ac4C | ac4C | Target | 286.1 | Wide | 154 | 5.106 | 1 | 85 | 9 |
| ac4C SILIS | ac4C SILIS | ISTD | 300 | Wide | 163 | 5.104 | 1 | 85 | 9 |
| acp3U | acp3U | Target | 346.1 | Wide | 214.1 | 1.963 | 1 | 95 | 15 |
| Am | Am | Target | 282.1 | Wide | 136 | 6.057 | 1 | 130 | 17 |
| Am SILIS | Am SILIS | ISTD | 298 | Wide | 146 | 6.055 | 1 | 130 | 17 |
| C | C | Target | 244.1 | Wide | 112 | 1.81 | 1 | 175 | 13 |
| C SILIS | C SILIS | ISTD | 256 | Wide | 119 | 1.806 | 1 | 175 | 13 |
| cm5U | cm5U | Target | 303.1 | Wide | 171.1 | 1.871 | 1 | 100 | 7 |
| Cm | Cm | Target | 258.1 | Wide | 112 | 3.845 | 1 | 180 | 9 |
| Cm SILIS | Cm SILIS | ISTD | 271 | Wide | 119 | 3.841 | 1 | 180 | 9 |
| D | D | Target | 247.1 | Wide | 115 | 1.621 | 1 | 70 | 5 |
| D SILIS | D SILIS | ISTD | 258 | Wide | 121 | 1.618 | 1 | 70 | 5 |
| G | G | Target | 284.1 | Wide | 152 | 4.144 | 1 | 130 | 17 |
| G SILIS | G SILIS | ISTD | 299 | Wide | 162 | 4.143 | 1 | 130 | 17 |
| GalQ | GalQ | Target | 572.3 | Wide | 295.5 | 4.069 | 1 | 115 | 20 |
| Gm | Gm | Target | 298.1 | Wide | 152 | 4.929 | 1 | 100 | 9 |
| Gm SILIS | Gm SILIS | ISTD | 314 | Wide | 162 | 4.926 | 1 | 100 | 9 |
| i6A | i6A | Target | 336.3 | Wide | 204.1 | 7.959 | 1 | 140 | 17 |
| i6A SILIS | i6A SILIS | ISTD | 356 | Wide | 219 | 7.953 | 1 | 140 | 17 |
| I | I | Target | 269.1 | Wide | 137 | 4.007 | 1 | 100 | 10 |
| I SILIS | I SILIS | ISTD | 283 | Wide | 146 | 4.004 | 1 | 100 | 10 |
| m1A | m1A | Target | 282.1 | Wide | 150 | 1.709 | 1 | 150 | 25 |
| m1A SILIS | m1A SILIS | ISTD | 298 | Wide | 161 | 1.719 | 1 | 150 | 25 |

| | | | | | | | | | |
|---|---|---|---|---|---|---|---|---|---|
| m1G | m1G | Target | 298.1 | Wide | 166 | 4.858 | 1 | 105 | 13 |
| m1G SILIS | m1G SILIS | ISTD | 314 | Wide | 177 | 4.856 | 1 | 105 | 13 |
| m2G | m2G | Target | 298.1 | Wide | 166 | 4.858 | 1 | 95 | 17 |
| m2G SILIS | m2G SILIS | ISTD | 314 | Wide | 177 | 4.856 | 1 | 95 | 17 |
| m3C | m3C | Target | 258.1 | Wide | 126 | 1.676 | 1 | 88 | 14 |
| m3C SILIS | m3C SILIS | ISTD | 271 | Wide | 134 | 1.675 | 1 | 88 | 14 |
| m3U | m3U | Target | 259.1 | Wide | 127 | 4.255 | 1 | 75 | 9 |
| m5C | m5C | Target | 258.1 | Wide | 126 | 3.599 | 1 | 185 | 13 |
| m5C SILIS | m5C SILIS | ISTD | 271 | Wide | 134 | 3.594 | 1 | 185 | 13 |
| m5U | m5U | Target | 259.1 | Wide | 127 | 4.255 | 1 | 95 | 9 |
| m5U SILIS | m5U SILIS | ISTD | 271 | Wide | 134 | 4.251 | 1 | 95 | 9 |
| m5Um | m5Um | Target | 273.2 | Wide | 127 | 5.613 | 1 | 90 | 13 |
| m6A | m6A | Target | 282.1 | Wide | 150 | 6.514 | 1 | 125 | 17 |
| m6A SILIS | m6A SILIS | ISTD | 298 | Wide | 161 | 6.509 | 1 | 125 | 17 |
| m6Am | m6Am | Target | 296 | Wide | 150 | 6.931 | 1 | 125 | 17 |
| m7G | m7G | Target | 298.1 | Wide | 166 | 3.17 | 1 | 100 | 13 |
| m7G SILIS | m7G SILIS | ISTD | 314 | Wide | 177 | 3.191 | 1 | 100 | 13 |
| m22G | m22G | Target | 312.1 | Wide | 180 | 5.691 | 1 | 105 | 13 |
| m22G SILIS | m22G SILIS | ISTD | 329 | Wide | 192 | 5.689 | 1 | 105 | 13 |
| m66A | m66A | Target | 296 | Wide | 164 | 7.141 | 1 | 130 | 21 |
| m66A SILIS | m66A SILIS | ISTD | 313 | Wide | 176 | 7.135 | 1 | 130 | 21 |
| m66Am | m66Am | Target | 310 | Wide | 164 | 7.423 | 1 | 120 | 15 |
| ManQ | ManQ | Target | 572.3 | Wide | 295.5 | 4.069 | 1 | 120 | 20 |
| mcm5s2U | mcm5s2U | Target | 333.1 | Wide | 201 | 6.345 | 1 | 92 | 8 |
| mcm5s2U SILIS | mcm5s2U SILIS | ISTD | 347.1 | Wide | 210 | 6.332 | 1 | 92 | 8 |
| mcm5U | mcm5U | Target | 317.1 | Wide | 185.1 | 4.996 | 1 | 95 | 5 |
| mcm5U SILIS | mcm5U SILIS | ISTD | 331 | Wide | 194 | 4.995 | 1 | 95 | 5 |

| | | | | | | | | | |
|---|---|---|---|---|---|---|---|---|---|
| ncm5s2U | ncm5s2U | Target | 318.1 | Wide | 186 | 4.071 | 1 | 95 | 7 |
| ncm5U | ncm5U | Target | 302 | Wide | 170 | 2.2 | 1 | 85 | 8 |
| ncm5U SILIS | ncm5U SILIS | ISTD | 316 | Wide | 179 | 2.21 | 1 | 85 | 8 |
| Q | Q | Target | 410.2 | Wide | 295.1 | 3.989 | 1 | 115 | 12 |
| s2U | s2U | Target | 261.1 | Wide | 129 | 4.535 | 1 | 80 | 6 |
| t6A | t6A | Target | 413.1 | Wide | 281.1 | 5.921 | 1 | 130 | 9 |
| t6A SILIS | t6A SILIS | ISTD | 434 | Wide | 297 | 5.92 | 1 | 130 | 9 |
| U | U | Target | 245.1 | Unit | 113 | 2.805 | 1 | 95 | 5 |
| U SILIS | U SILIS | ISTD | 256 | Unit | 119 | 2.826 | 1 | 95 | 5 |
| Um | Um | Target | 259.2 | Wide | 113 | 4.559 | 1 | 96 | 8 |
| Um SILIS | Um SILIS | ISTD | 271.1 | Wide | 119 | 3.841 | 1 | 96 | 8 |
| Y | Y | Target | 245.1 | Wide | 209 | 1.68 | 1 | 90 | 5 |
| Y SILIS | Y SILIS | ISTD | 256 | Wide | 220 | 1.678 | 1 | 90 | 5 |
